# An RNA binding module of SWI/SNF is required for activation of cell-type specific enhancers and super-enhancers in early development

**DOI:** 10.1101/2022.09.06.506785

**Authors:** Dhurjhoti Saha, Srinivas Animireddy, Junwoo Lee, Yuan-chi Lin, Kyle Feola, Abhinav K Jain, Yue Lu, Bin Liu, Blaine Bartholomew

## Abstract

The mammalian SWI/SNF complex is an ATP-dependent chromatin remodeler and master regulator in development that when mutated is the cause for several human diseases including cancer. Although SWI/SNF is highly enriched at enhancers and its basic chromatin remodeling activities have been studied for over 30 years, there is little known about how it regulates enhancer activity or enhancer-promoter interactions. We find a putative RNA binding module located near the C-terminus of the catalytic subunit of SWI/SNF required for SWI/SNF recruitment to cell-type specific enhancers and super-enhancers in naïve and cell lineage primed pluripotent cells. The AT-hook is required for acquisition of the active histone marks H3K27ac and H3K4me1 and recruitment of the MLL3/4 co-activator to these enhancers and super-enhancers. Consistent with changes in enhancer architecture, loss of the AT-hook interferes with activation of genes involved in cell lineage priming as well as genes normally activated in naïve pluripotent cells.

## Introduction

The SWI/SNF chromatin remodeler is a master epigenetic regulator that controls cell fate determination and embryonic stem cell proliferation and maintenance (Ho and Crabtree, 2010; Lessard and Crabtree, 2010). Central in this regulation are distal regulatory elements called enhancers that act at great distances from their target genes and facilitate lineage and cell-type specific transcription (Calo and Wysocka, 2013). Of the three classes of SWI/SNF complexes, BAF, PBAF and GBAF or ncBAF, only the BAF complex is preferentially bound to enhancers (Gatchalian et al., 2018; Lessard et al., 2007; Michel et al., 2018; Park et al., 2021). BRG1/SMARCA4, one of two catalytic subunits of the BAF complex and which we will refer to as BRG1 from here on out, has been shown to be recruited to various enhancers by sequence-specific DNA binding transcription factors (TFs) (Trotter and Archer, 2008; Wood et al., 2016; Yu et al., 2013). BRG1 also promotes TF binding to enhancers often in conjunction with increasing accessibility at the enhancers (Bossen et al., 2015; Hu et al., 2011; Ni et al., 2008).

Recruitment of BRG1 however seems to involve more than just TFs as seen by BRG1 requiring other factors for its recruitment. P300 is a histone acetyltransferase that acetylates lysine 27 of histone H3 (H3K27ac) at enhancers and is a hallmark of active enhancers. Enhancers and super-enhancers are dynamically activated by H3K27 acetylation which promotes TFIID and RNA polymerase (RNAP)II recruitment to nearly all enhancers (Narita et al., 2021). BRG1 physically interacts with p300, together they facilitate each other’s binding, and BRG1 promotes robust acetylation of H3K27 to activate cell-type specific enhancers (Alexander et al., 2015; Alver et al., 2017; Blumli et al., 2021). P300 has been observed to bind 122 lncRNAs and for lncSmad7 to facilitate p300 recruitment to enhancers (Maldotti et al., 2022). CBP, the paralog of p300, has also been shown to interact with enhancer (e)RNAs and eRNA to activate its catalytic activity (Bose et al., 2017). The other co-activator that promotes BRG1 binding is MLL3/4, a histone methyltransferase that monomethylates lysine 4 on histone H3 (H3K4) at enhancers and with BRG1 reciprocally promote each other’s binding to enhancers involved in adipogenesis (Park et al., 2021). MLL3/4 is also required for p300/CBP binding at enhancers and for the formation of super-enhancers (Lai et al., 2017). MLL3/4 promotes long-range chromatin interactions and facilitates those between enhancers and promoters (Yan et al., 2018). Some of the outstanding questions is how these 3 factors work together for enhancer activation and if RNA might play a part in the three-way co-dependency of BRG1, p300 and MLL3/4 recruitment.

Evidence is beginning to accumulate that RNA can facilitate the recruitment of SWI/SNF in addition to the protein factors described earlier, similar to that observed for CTCF and PRC2 (Brockdorff, 2013; Hansen et al., 2019; Saldana-Meyer et al., 2019; Yan et al., 2019). The long non-coding (lnc)RNA *lncTC7* promotes BAF binding to the *TCF7* locus; lncRNA *lincRNA-Cox2, IL-7-AS* and *MALAT1* target BAF to proinflammatory response genes (Hu et al., 2016; Huang et al., 2019; Liu et al., 2019; Wang et al., 2015) and *SWINGN* promotes SWI/SNF binding through the SMARCB1 subunit (Grossi et al., 2020). RNAs can also compete BAF complexes away from DNA or inhibit their remodeling activity(Cajigas et al., 2015). Some of the subunits or domains implicated to interact with SWI/SNF are SMARCB1, BRD4 and BRG1 subunits and within BRG1 there is evidence the HSA/BRK, helicase and bromo domains bind RNA (Patty and Hainer, 2020). Most of the interactions however with regions of BRG1 so far have been shown to negatively regulate its activity or compete it away from DNA.

In BRG1 between the SnAC and bromo domains is an AT-hook motif which at its center is an arginine-glycine-arginine flanked on one or both sides by proline that binds preferentially to the minor groove of AT-rich DNA (Bewley et al., 1998; Huth et al., 1997). Other AT-hooks like that found in BRG1 have extended GGR repeats or additional basic residues farther from the core that cause the AT-hook to have a higher affinity for RNA than for DNA (Dickinson and Kohwi-Shigematsu, 1995; Filarsky et al., 2015; Sears et al., 2004). The AT-hook domain is found in a variety of proteins with most of them being chromatin modifiers and the AT-hook has been suggested to anchor these proteins to chromatin (Aravind and Landsman, 1998; Filarsky et al., 2015). The AT-hook is required for NURF and RSC remodeling, but its role in remodeling and potential target(s) in these cases are not known (Cairns et al., 1999; Xiao et al., 2001). It remains to be determined if the AT-hook in the catalytic subunit of SWI/SNF regulates the remodeling activity of SWI/SNF or is required for stable binding of SWI/SNF to chromatin. We have found the AT-hook of BRG1 in mouse embryonic stem cells is vital for BRG1 recruitment to many cell-type specific enhancers and super-enhancers and in turn regulates the recruitment of MLL3/4 and the acquisition of H3K27ac and H3K4me1.

## Results

### The AT-hook of BRG1 is required for BRG1 recruitment at many but not all targets

We investigated the AT-hook of BRG1 by first deleting exon 33 containing the AT-hook in both copies of BRG1 in mouse embryonic stem cells (mESCs) and refer to them as dAT. The genome wide binding patterns of WT and two independent clones of the dAT mutant (dAT1 and dAT2) BRG1 were mapped in two distinct pluripotent states referred to as naïve and primed, representative of the pre- and post-implantation stages, using BRG1 ChIP-seq and CUT&RUN. We observed in these two different cell types that most of the sites where BRG1 binds are cell-type specific with significantly less that are shared between naïve and primed (Figures 1A-B and S1A). BRG1 in mESCs is assembled into esBAF and GBAF or ncBAF complexes, and the esBAF complex preferentially binds to cis-regulatory regions and the GBAF complex to promoter regions (Alpsoy and Dykhuizen, 2018; Gatchalian et al., 2018). About 80% of the naïve- and primed-specific BRG1 peaks were at intronic or intergenic regions and most likely correspond to the esBAF complex and only about 6-13% of the peaks associate with promoter regions (Figure 1C). CUT&RUN is more prone than ChIP-seq to detect BRG1 binding in naïve cells with 39,915 naïve specific peaks detected by CUT&RUN versus only 2,825 naïve specific peaks detected by ChIP-seq (Figures 1A-B and S1A). BRG1 binding is reduced at half of its sites using ChIP-seq in the primed state when the AT-hook is deleted as compared to 82% of the BRG1 sites being reduced or lost when detected by CUT&RUN (Figures 1D-E and S1C-D). Slightly different effects are seen at naïve binding sites of BRG1 with 86% of the binding sites lost when the AT-hook is deleted as detected by ChIP-seq and 49% of the binding sites lost when detected by CUT&RUN (Figures 1E and S1C-D). Overall, there is good correlation between the two methods for mapping the genome-wide binding of BRG1 with 25-30% of the ChIP-seq peaks overlapping with those detected by CUT&RUN (Figure S1B). Deletion of the AT-hook also causes a shift in BRG1 positioning with some pre-existing BRG1 sites gaining BRG1 with loss of the AT-hook (Figures S1E-F). In summary, we find the AT-hook has a major role in BRG1 recruitment genome-wide and also is likely not the only mode of BRG1 recruitment.

**Figure 1.**
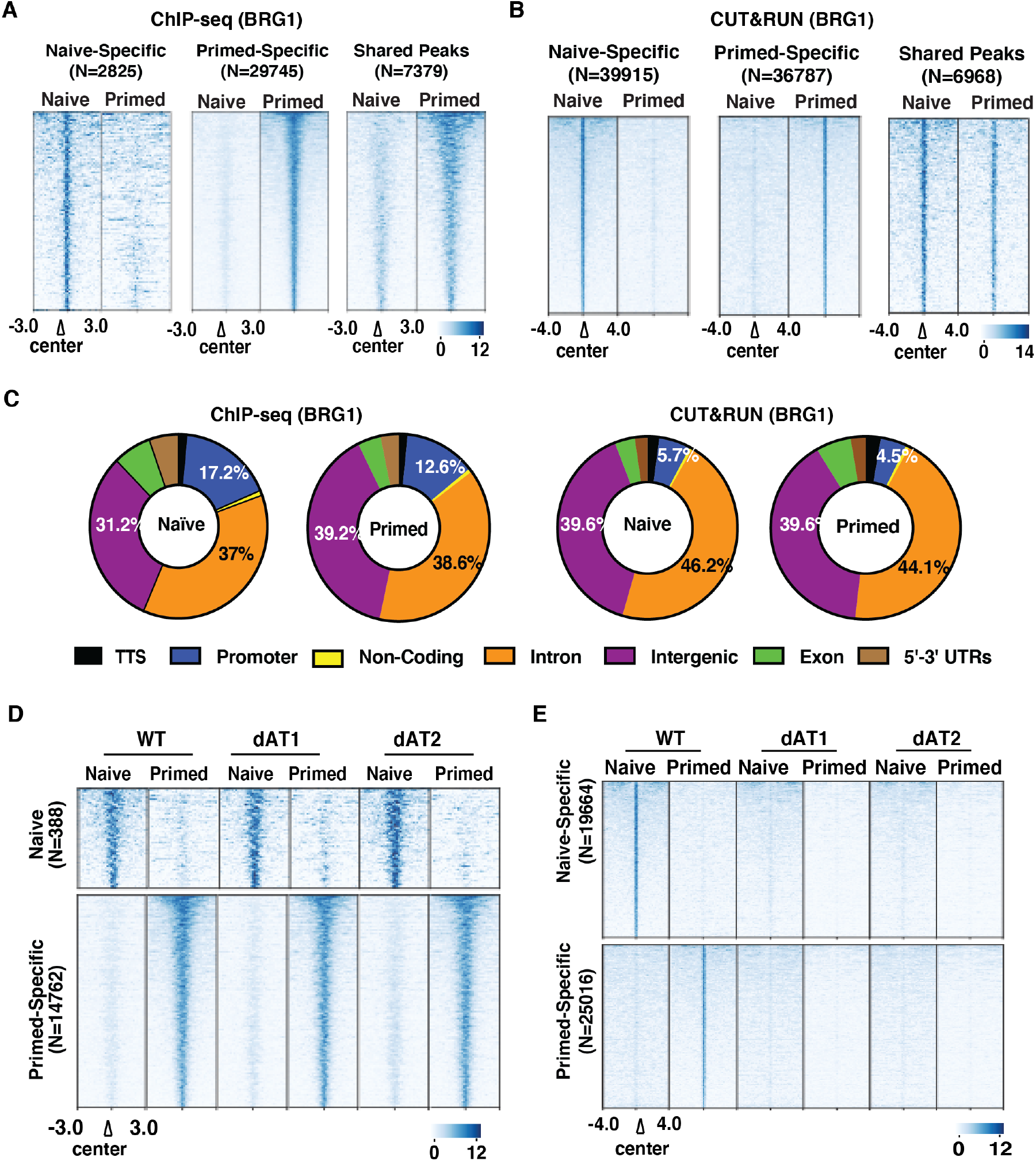
Two orthogonal approaches show the AT-hook of BRG1 facilitates BRG1 recruitment in naïve and primed states. (A-B) Heatmaps show BRG1 localization using (A) ChIP-seq or (B) CUT&RUN in wild type (WT) and two AT-hook deletion mutants (dAT1, dAT2) that are unique to naïve and primed cells or in common for both. (C) Pie charts show the genome-wide distribution of BRG1 using ChIP-seq (left) and CUT&RUN (right). (D) The localization of a significant number of BRG1 sites detected by ChIP-seq are not affected by loss of the AT-hook of BRG1. (E) In contrast a large group of BRG1 binding sites detected by CUT&RUN are lost when the AT-hook is absent. Signals are sorted based on WT in each group; N represents the total number of peaks.

### The co-dependent recruitment of BRG1 and acquisition of H3K27ac at enhancers is AT-hook dependent

We investigate the interplay between the AT-hook and other factors known to be involved in BRG1 recruitment starting with p300/CBP, a co-activator that is principally responsible for acetylation of lysine 27 of histone H3 (H3K27ac) at enhancers and focus on where H3K27ac and BRG1 occupy adjoining sites. BRG1 physically interacts with p300/CBP and they are co-dependent for their recruitment with BRG1 also positively regulating p300 acetylation activity (Manickavinayaham et al., 2019; Naidu et al., 2009; Wu et al., 2018; Yang et al., 2019). In the naïve and primed states, the number of BRG1 sites that co-localize with H3K27ac are respectively 94% and 46% of all BRG1 binding sites in cis-regulatory regions (Figure S2A). When the AT-hook is deleted, we observe that both H3K27ac and BRG1 binding are reduced at these regions in naïve and primed cells (Figure 2A). These data show that p300/CBP cannot compensate for the loss of the AT-hook for BRG1 recruitment and implies that binding of the AT-hook to RNA might be the step that initiates the co-recruitment of BRG1 and p300/CBP or for BRG1 to stimulate the enzymatic activity of p300/CBP.

**Figure 2.**
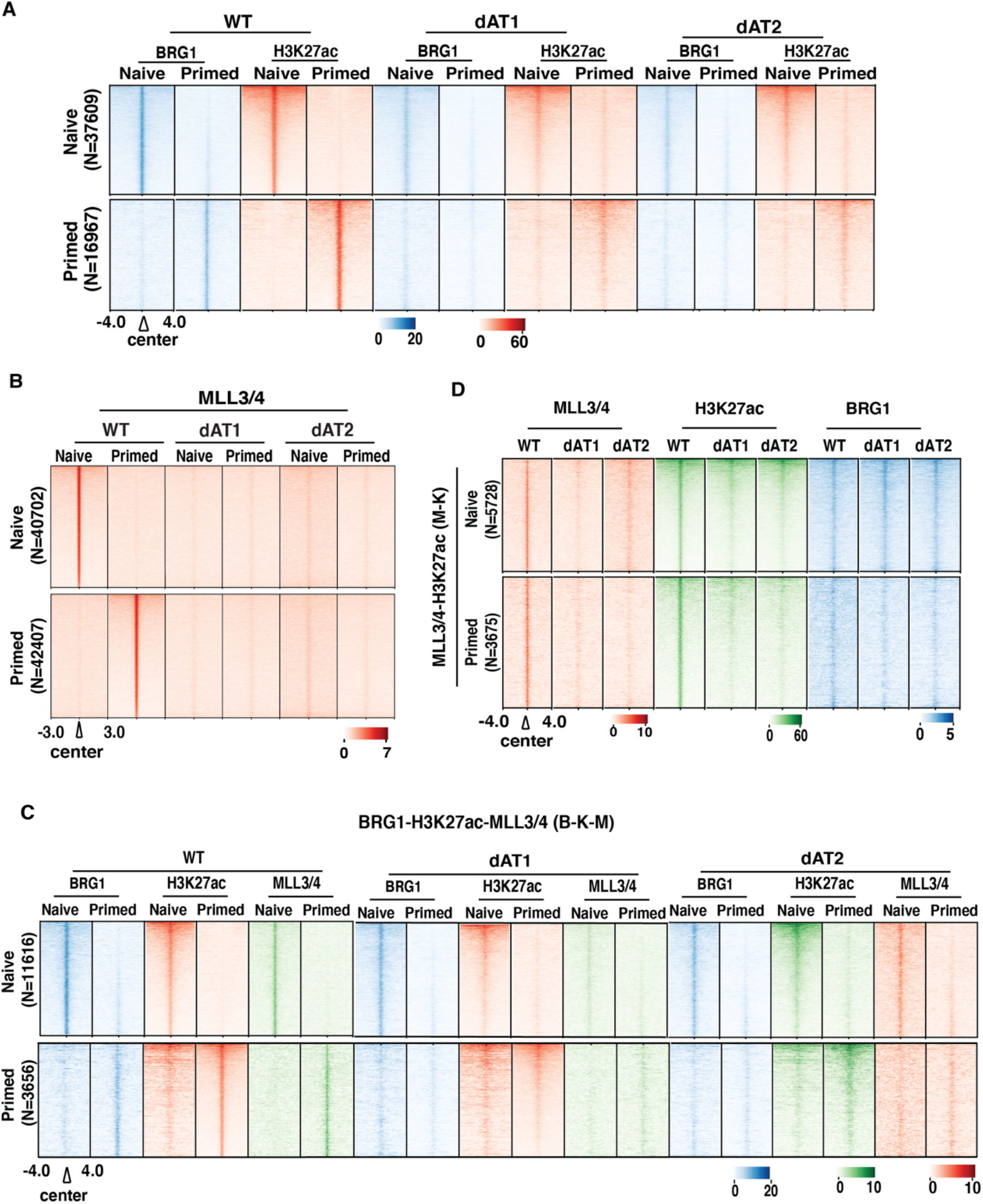
BRG1 and its AT-hook are required for acquisition of H3K27ac and recruitment of MLL3/4. (A) The heatmap shows the regions where BRG1 (blue) and H3K27ac (red) overlap in a cell-type specific manner in naïve (upper panel) and primed (lower panel) cells for wild type (WT) and two independent clones that lack the AT-hook (dAT1 and dAT2). (B) MLL3/4 localization at cis-regions that are AT-hook dependent in navie and primed states. (C-D) The localization of BRG1(blue), H3K27ac (red), and MLL3/4 (green) in wild type and AT-hook deletion mutants at BRG1, MLL3/4 and H3K27ac co-localized (C) and MLL3/4-H3K27ac colocalized (D) in naïve and primed states; number of sites corresponds to those sets shown in Figure S2J.

### Recruitment of the MLL3/4 complex requires BRG1 and it AT-hook

We expand our investigation into BRG1 recruitment by examining MLL3/4, another co-activator that has been shown to mediate BRG1 recruitment at cis-regulatory elements in adipogenesis (Park et al., 2021). As detected by CUT&RUN, MLL3/4 binds at sites exclusively in the naïve or primed state that are primarily intergenic and intronic regions rather than promoters (Figures S2B and S2D). MLL3/4 binding at the intergenic and intronic regions is highly dependent on the AT-hook of BRG1, whereas most of the sites where MLL3/4 is bound in both states do not depend on the AT-hook for their recruitment (Figures 2B and S2C, S2E-F). These data indicate BRG1 and its AT-hook are preferentially required for stage-specific recruitment of MLL3/4 to cis-regulatory regions. In contrast to the co-dependency of BRG1 and H3K27ac observed earlier, many of the sites where MLL3/4 binding depends on the AT-hook of BRG1 are sites where BRG1 enrichment is relatively low compared to that observed where BRG1 and H3K27ac are co-localized (compare Figure 2A to S2G). There are respectively 20% and 30% of the AT-hook dependent MLL3/4 sites in naïve and primed that are however significantly enriched with BRG1 and BRG1 recruitment at these sites is AT-hook dependent (Figures S2H-I). These data are consistent with two types of MLL3/4 sites that depend on the AT-hook of BRG1 for their recruitment. Those sites with high BRG1 occupancy are ones where there is a reciprocal binding dependency; whereas the low BRG1 occupancy MLL3/4 sites are those that are probably more dependent on the catalytic activity changes in BRG1 caused by deletion of the AT-hook than direct physical interaction.

We examined the potential 3-way synergy between BRG1, H3K27ac as an indicator of p300/CBP, and MLL3/4 and divided them into two groups based on the level of BRG1 enrichment. In the highly enriched BRG1 group, we observe BRG1, H3K27ac and MLL3/4 enriched at naïve or primed-specific cis-regulatory regions (Figures 2C and S2J). At these sites deletion of the AT-hook causes the loss of all three in naïve or primed cells and highlights that MLL3/4 and p300/CBP combined are not able to offset the loss of BRG1 or themselves when the AT-hook of BRG1 is deleted. In the group of low BRG1 enrichment, there is loss of H3K27ac and MLL3/4 when the AT-hook is deleted but BRG1 binding is not altered (Figure 2D). These data are consistent as mentioned earlier with two distinct modes of BRG1 mediating the acquisition of H3K27ac and recruitment of MLL3/4 and highlights the central role of the AT-hook in both.

### Monomethylation of lysine 4 of histone H3 depends on the AT-hook of BRG1, but not in conjunction with stable MLL3/4 binding

The MLL3/4 catalytic subunit is primarily responsible for monomethylation of lysine 4 of histone H3 (H3K4me1) in mESCs (Dorighi et al., 2017; Hu et al., 2013) and prompted our investigation of the relationship between MLL3/4, H3K4me1 and the AT-hook of BRG1. About 80% of the H3K4me1 peaks in CUT&RUN are at intronic and intergenic sites. Many of the H3K4me1 intronic and intergenic sites are found in both states and a lesser number of sites are naïve- or primed-specific (Figure S3A). The cell-type specific H3K4me1 sites depend on the AT-hook of BRG1 for their proper localization, consistent with the AT-hook dependency of MLL3/4 (Figures 3A-B and S3B). There is also a smaller subset of H3K4me1 sites that do not depend on the AT-hook of BRG1 (Figures S3C-D). Enrichment of MLL3/4 and BRG1 is low at the AT-hook dependent H3K4me1 cis-regulatory regions, and both their recruitment is modestly dependent on the AT-hook of BRG1 (Figures S3E-F). We look next at the colocalization of MLL3/4 and H3K4me1 in the naïve and primed states and find at least half of the highly enriched MLL3/4 sites don’t colocalize with H3K4me1 (Figures 3C-D), consistent with that previously observed in mESCs (Dorighi et al., 2017). At sites where both MLL3/4 and H3K4me1 are enriched, H3K4me1 is not dependent on the AT-hook of BRG1 even though MLL3/4 recruitment is highly dependent on the AT-hook (Figures 3C-D). We first conclude from this data that the sites where MLL3/4 is most enriched or potentially the sites where its binding is most stabilized do not correspond to sites where H3K4me1 is likely to be acquired, consistent with prior observations of MLL3/4 facilitating transcription and eRNA synthesis independent of its catalytic activity (Dorighi et al., 2017). There also appears to be two types of H3K4me1 sites that vary based on their AT-hook dependency and levels of MLL3/4 co-occupancy. Only when MLL3/4 occupancy is low does monomethylation of H3K4me1 require the AT-hook of BRG1. Although this data could be interpreted as MLL3/4 not being the enzyme responsible for H3K4me1 at cis-regulatory regions, we don’t think this is the case based on prior data with the catalytically dead version of MLL3/4 (Dorighi et al., 2017). It seems more likely that the rapid turnover of MLL3/4 is vital for its enzymatic activity and monomethylation of H3K4 rather than the stable binding of MLL3/4.

**Figure 3.**
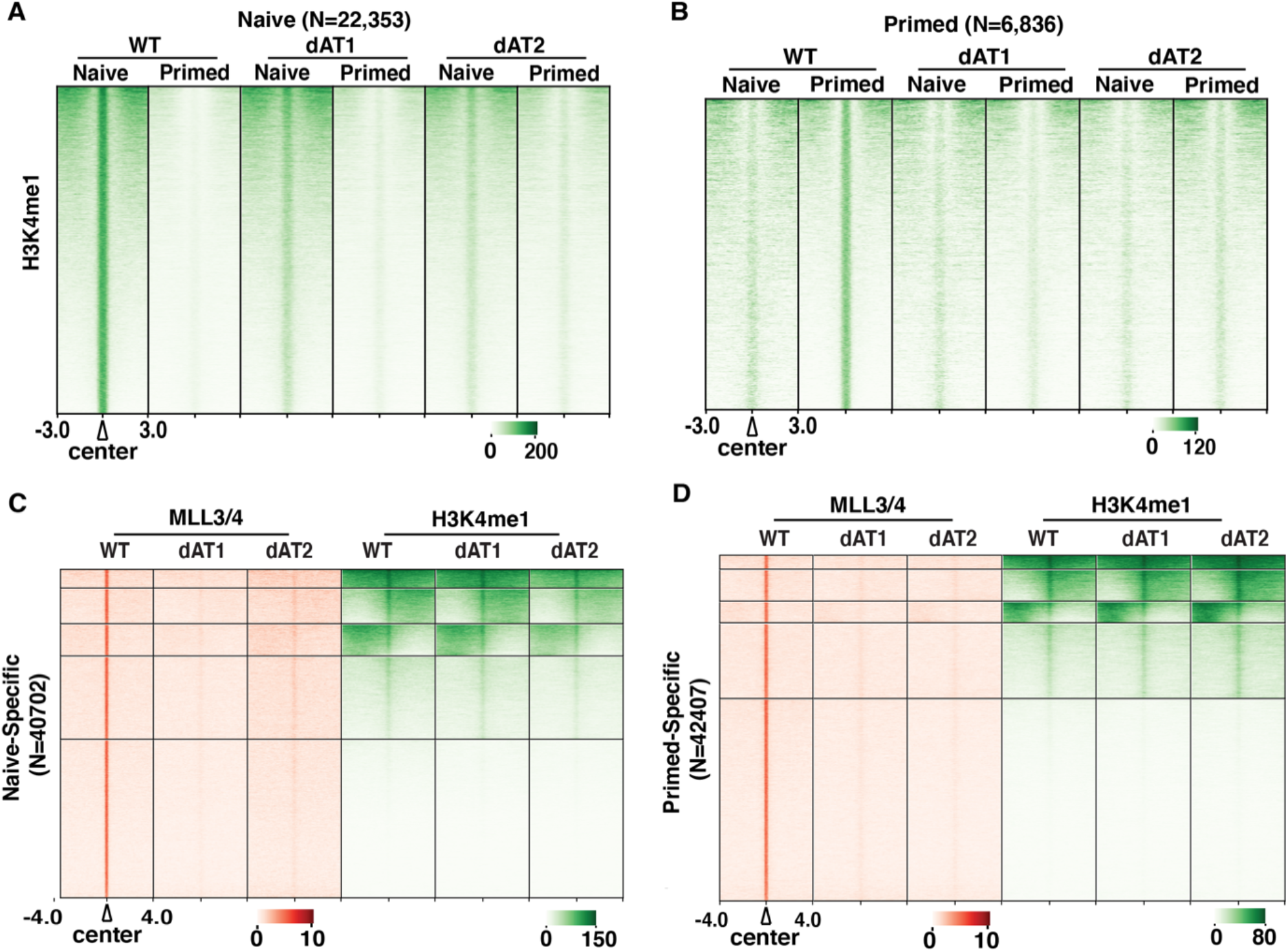
Monomethylation of H3K4 requires BRG1 and its AT-hook at a subset of its targets. (A-B) Heatmaps of all H3K4me1 (green) peaks detected by CUT&RUN for wild type and AT hook deletion mutants in naïve (A) and primed (B) cells are shown. (C-D) The heatmaps shows the extent which MLL3/4 (red) and H3K4me1 (green) overlap in naïve (C) and primed (D) cells in cis-regulatory regions and their dependency on the AT-hook of BRG1. (E) Heatmaps of H3K4me1 (green), H3K27ac (red) and BRG1 (blue) are shown for those regions where BRG1 and H3K27ac co-localize in only naïve or primed cells.

### BRG1 and its AT-hook is required for sequence-specific DNA binding transcription factor recruitment

BRG1 recruitment is also mediated by DNA sequence-specific binding transcription factors (TFs) such as nuclear receptors and acidic activation type factors(Hu et al., 2011; Huang et al., 2002; Laurette et al., 2015; Trotter and Archer, 2008). The role of BRG1 in TF recruitment is investigated using ATAC-seq and motif analysis of the protected DNA sequence to reveal the TFs bound proximal to BRG1 (Bentsen et al., 2020). There are a total 38,416 and 42,303 ATAC accessible peaks detected in naïve and primed cells and of these peaks there are 18,814 peaks in common between naïve and primed (Figure S4A). About 14,000 peaks reside in the intronic and intergenic regions that are either naïve- or primed-specific (Figure S4A). BRG1 ChIP-seq was used to examine accessible sites detected by ATAC-seq proximal to BRG1 as they showed a significantly higher number of BRG1 peaks proximal to ATAC peaks than with CUT&RUN. Over ¼ (27%) of all the primed BRG1 binding sites detected by ChIP-seq are proximal to accessible sites compared to only 5% of BRG1 peaks detected by CUT&RUN. The 7,711 primed-specific BRG1 peaks proximal to these accessible sites had no loss of BRG1 binding when the AT-hook was deleted; however, 80% of the other ¾ of the BRG1 primed-specific peaks (20,759) had significantly reduced binding when the AT hook was deleted (compare Figures 4A and 4B). These data suggest TF-mediated recruitment of BRG1 stabilizes the chromatin interactions of BRG1, thereby compensating for the loss of the AT-hook. Many of the regions where TF binding sites are detected by ATAC-seq appear to be active enhancers based on the co-localization of histone marks H3K27ac and H3K4me1 (Figures 4C and S4B). Primed-specific enhancers have elevated levels of BRG1, consistent with BRG1 being preferentially enriched at enhancers (Alexander et al., 2015; Bossen et al., 2015); whereas naïve-specific enhancers have BRG1 that is not exclusive to naïve cells (Figure 2C). Motif analysis using HOMER revealed the naïve ATAC peaks represent the binding of pluripotency TFs like Oct4, Nanog, Sox2 and members of the KLF TF family; whereas the primed ATAC peaks corresponded to epiblast specific TFs like Zic2, Zic3, Otx2 and Glis3 (Figures S5C-D). Even though they have started the transition to cell fate determination, primed cells have sites where Oct4, Sox2, Nanog and Tcf TFs are bound and is expected given these cells remain pluripotent.

**Figure 4.**
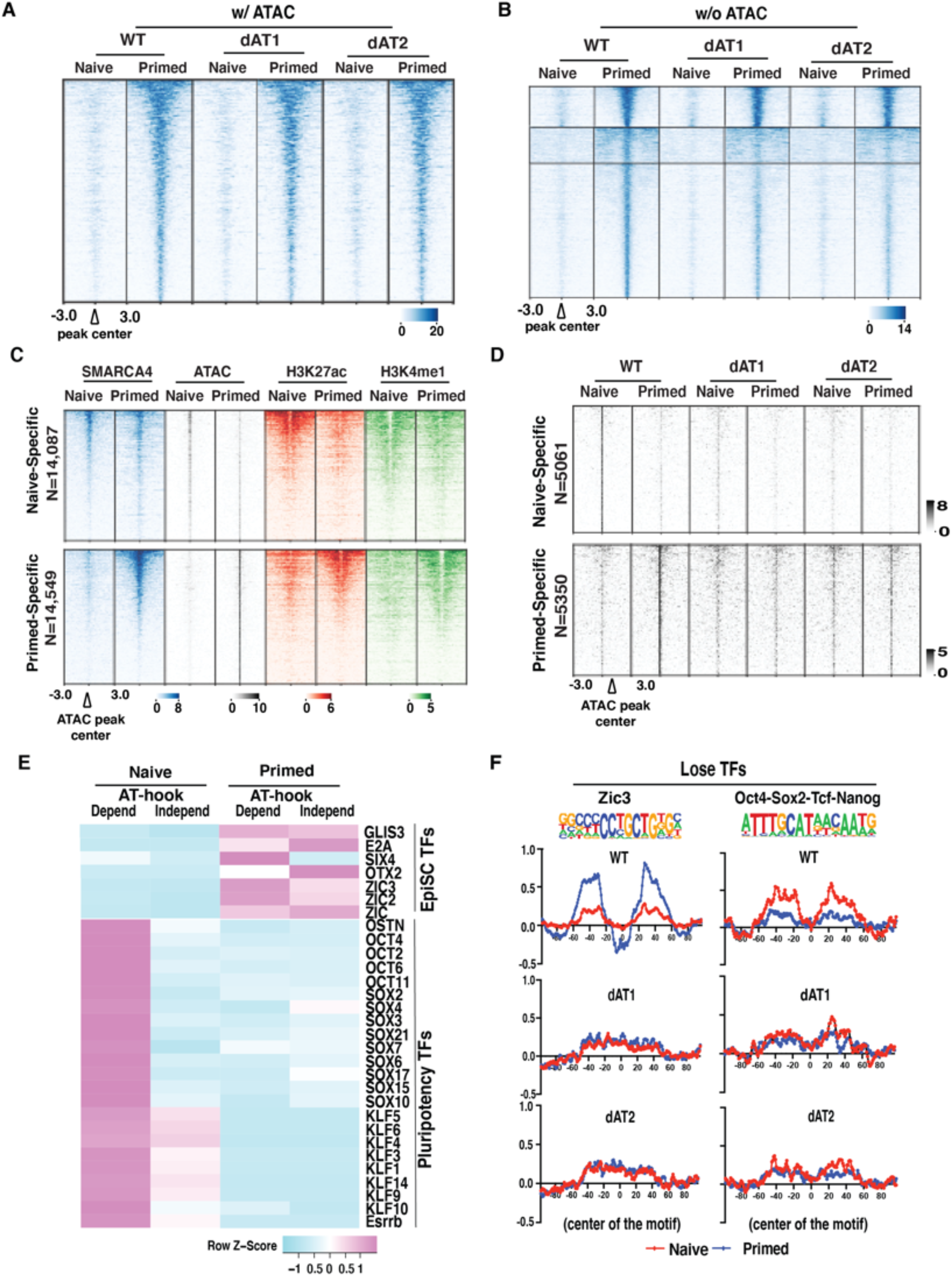
AT-hook of BRG1 is required for stage-specific TF recruitment. (A) Venn diagram shows naïve and primed specific intronic-intergenic ATAC-seq peak. (B) Heatmap showing SMARCA4 localization (blue), ATAC-signals (grey), and active enhancer histone marks (H3K27ac [red], and H3K4me1 [green]) at naïve (top) and primed (bottom) specific intronic-intergenic ATAC-seq peaks. ChIP-seq signals are sorted based on WT SMARCA4 in each condition. (C) Heatmap showing ATAC signals in the two independent clones of the AT-hook mutant in naïve (top) and primed (bottom) states versus WT. ATAC signals are sorted based on WT. (D) Heatmap shows the ATAC-signals for WT and dAT-mutants in naïve (top) and primed (bottom) conditions that are unaltered; N represents the number of intronic-intergenic ATAC-seq peaks in each category. (E) Heatmap showing the pluripotency and epiblast specific (EpiSC) specific transcription factors (TFs) motif enrichment between AT-hook dependent -vs.-AT-hook independent ATAC-seq intronic-intergenic peaks in naïve and primed states. (F) DNA footprinting shows loss of primed/EpiSC specific TF Zic3 (left panel) and pluripotency TFs <u>O</u>ct4-<u>S</u>ox2-<u>T</u>cf-<u>N</u>anog (right panel) binding in the AT-hook deletion mutants.

Next, we examined if TF binding was dependent on the AT-hook of BRG1 and found 5,061 and 5,350 accessible sites in respectively the naïve and primed state that depend on the AT-hook of BRG1 (Figures 4D-E and S5A-B). BRG1 binding to these sites does not change when the AT-hook is deleted whether TF binding or accessibility is apparently retained (Figures S4C-D). These regions are further indicated to be active enhancers by the enrichment of histone modifications H3K4me1 and H3K27ac (Figures S4E-F). Footprint analysis illustrates how deletion of the AT-hook of BRG1 impacts the primed-specific binding of Zic3 and the naïve-specific binding of Oct4-Sox2-Tcf-Nanog (Figure 4F). There are also sites where Oct4-Sox2-Tcf-Nanog binds in both naïve and primed cells that are not affected by loss of the AT-hook as well as Otx2 binding in primed cells (Figures S5E-F). BRG1 had previously been shown to be required for recruitment of OCT4, SOX2 and NANOG when BRG1 is eliminated (Barisic et al., 2019; Ho et al., 2011; King and Klose, 2017; Miller et al., 2017), but this is the first time BRG1 has been shown to be vital for recruitment of cell lineage commitment TFs and for its AT-hook to be required for binding of both pluripotency- and epiblast-specific TFs.

### BRG1 and its AT-hook are required to activate transcription in the naïve and primed states

Given the extensive restructuring of enhancers that occurs in naïve and primed cells when the AT-hook of BRG1 is deleted, we examined changes in gene expression by mapping nascent transcription at near base-pair resolution using <u>P</u>recision <u>R</u>un-<u>O</u>n Sequencing (PRO-seq) in naïve and primed cells for coding and noncoding genes. While the number of genes dysregulated by deletion of the AT-hook varied between the two independent dAT clones the patterns and trends were similar (compare Figures 5 to S6). Consistent with enhancer switching that occurs between the naïve and primed states we observe a high number of genes that are primarily expressed in only naïve or primed cells. The ∼3600-4500 genes specifically upregulated in naïve are divided into groups based on how they respond to deletion of the AT-hook (Figures 5A and 5C and, S6A-B). One set of the naïve specific genes fails to be activated in the dAT mutants and the other is also down regulated but fails to be shut down when transitioning to primed. Gene onotology reveals that the group that fails to be repressed in primed include genes encoding factors important for pluripotency that likely need to be shut down as cells begin to exit pluripotency (Figure 5D) (Tsogtbaatar et al., 2020). These genes encode factors involved in the metabolic shift from naïve to primed that occurs with shut down of oxidative phosphorylation (OX-PHOS) and pentose phosphate pathways (PPP) accompanied by an increase in glycolysis (Stincone et al., 2015; Tsogtbaatar et al., 2020). Other signature events for the gradual decommissioning of the pluripotent state that we see in the transcriptional changes are disruptions in lipid metabolism (Cornacchia et al., 2019; Tanosaki et al., 2020; Wang et al., 2017), cell cycle control (Ter Huurne et al., 2017) and G protein-coupled receptor signaling (Callihan et al., 2011; Dolatshad et al., 2015).

**Figure 5.**
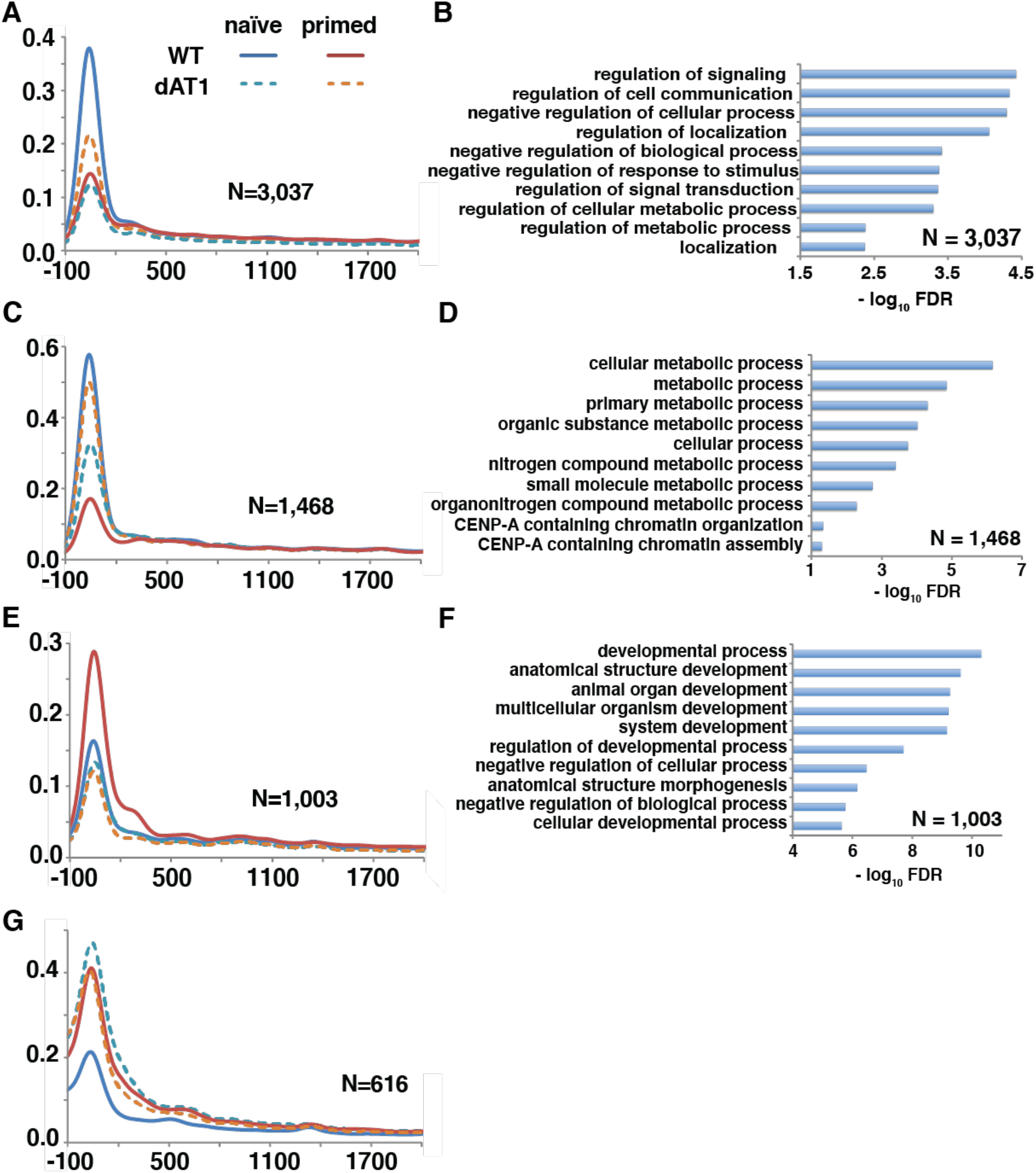
Loss of the AT-hook of SMARCA4/BRG1 causes transcription dysregulation in both the naïve and primed states. (A-D) Meta-analysis of PRO-seq signals (upstream TSS −100 bp to +300 bp downstream of TSS) shows differential pausing genes between WT and dAT1 mutant in naïve (A, B) and primed states (C, D). Paused Up-regulated genes in WT are shown in panel (A) and (C) and panel (B) and (D) show paused genes that are misregulated in dAT1 mutant in naïve (B) and primed (D) respectively (E-G) Bargraphs show gene ontology (GO) term enriched in WT pausing up-regulated genes in naïve and primed (group-I and group - III) and misregulated genes in dAT1 (group - II).

There are genes that also fail to be activated when the AT-hook of BRG1 is deleted that are specific to primed cells (Figures 5E and S6B). This group of genes are related to development, cell differentiation, and cell-cell adhesion; consistent with early cell lineage priming occurring in the primed state that require BRG1 and its AT-hook (Figure 5F). A second set of genes that are normally expressed primarily in primed cells are prematurely turned on in naïve cells upon loss of the AT-hook (Figure 5G). We next examined whether changes in chromatin structure at the promoters of the target genes could be responsible for their dysregulation when the AT-hook of SMARCA4/BRG1 is deleted. Mapping both DNA accessibility and enrichment of trimethylated lysine 4 of histone H3 (H3K4me3) at promoter regions showed no significant changes at genes where transcription is altered by loss of the AT-hook (Figures S7A-F). Together these data show BRG1 and its AT-hook is required for activation of both naïve- and primed-specific gene expression as expected by the changes we observe at naïve and specific enhancers. We have also observed that BRG1 has a dual role in repressing other genes and an important role in the gradual decommissioning of the pluripotent state.

### Cell-type specific super-enhancers in naïve and primed cells require BRG1 and its AT-hook

We investigate the role of BRG1 and its AT-hook in controlling cell identity by first mapping the genomic location of the MED1 subunit of the Mediator complex to identify cell type specific super-enhancers. MED1 binds TFs and potentially eRNAs, the majority of MED1 is an intrinsically disordered region spanning from amino acids 518-1581 and is likely involved in the recruitment of Mediator to transcription hubs or condensates (Chen et al., 2021; Hsieh et al., 2014; Klein et al., 2020). MED1 is found to preferentially bind to intronic and intergenic regions that are either cell-type specific or at sites present in both (Figures 6A-B). The initial super-enhancers are based on all naïve and primed MED1 binding sites being ranked by enrichment and clustering together and were sorted based on being present only in naïve and primed or in both (Figures 6C-D). These super-enhancers are clusters of individual enhancers that are regulated by BRG1. We observe within these super-enhancers sites that depend on BRG1 and its AT-hook for acquiring H3K27ac and MLL3/4 which overlap with 92-95% of the naïve and primed specific super-enhancers (Figure 6E). On the average there are multiple of these AT-hook dependent features per enhancer ranging from 3-9 per super-enhancer. These data show BRG1 and its AT-hook have an important role in activating super-enhancers by facilitating the cooperative recruitment of BRG1, MLL3/4 and H3K27ac. We also observe the naïve and primed-specific AT-hook regulated super-enhancers are within 500 kb of respectively approximately 60% and 30% of those genes that are differentially regulated in naïve and primed states when the AT-hook is deleted, consistent with these super-enhancers regulating transcription in a BRG1-dependent manner (Figure 6F).

**Figure 6.**
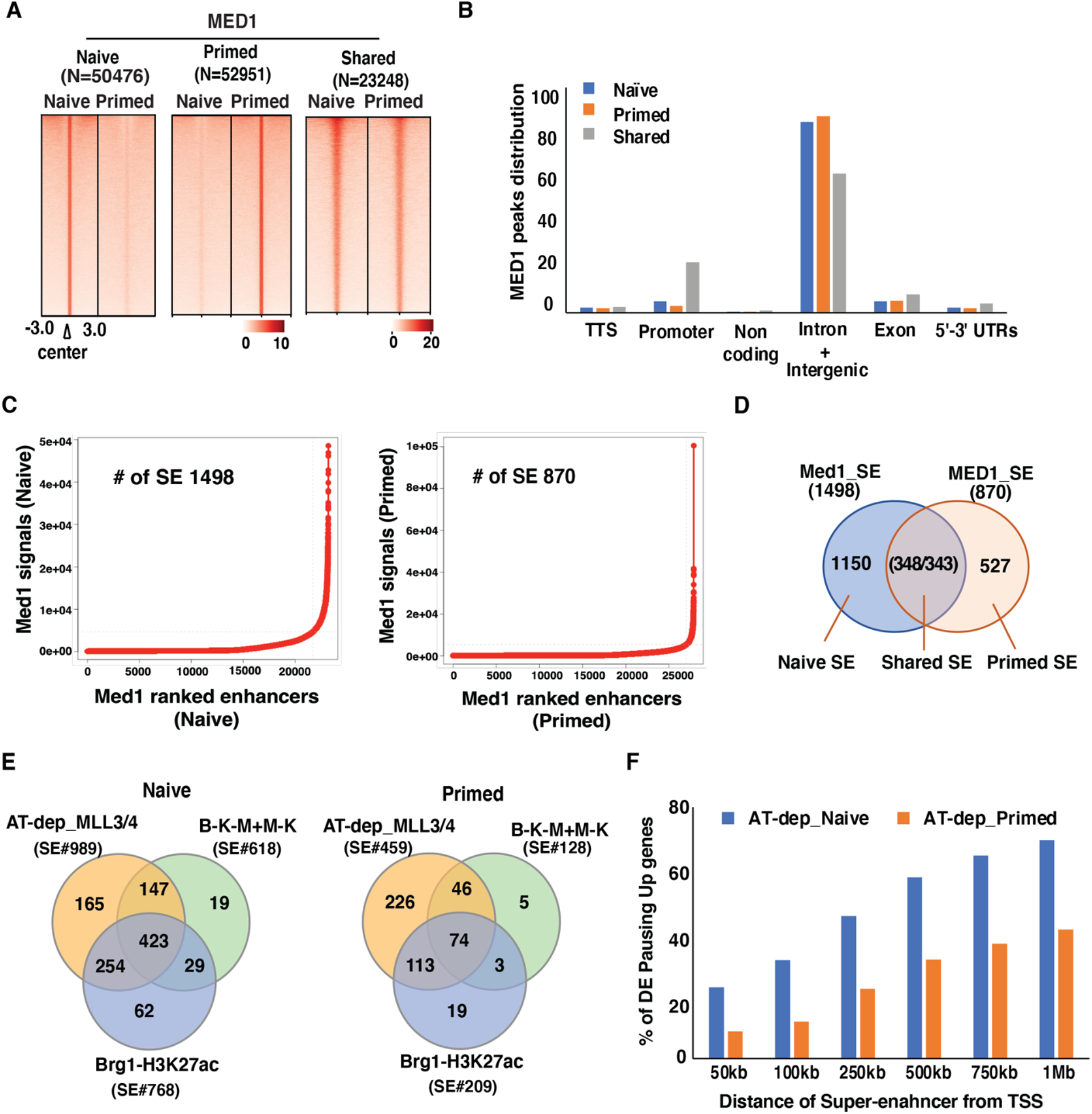
Activation of cell-type specific super-enhancers requires BRG1 and its AT hook. (A) The heatmaps show WT Med1 localization in naïve (left), primed (middle), and shared (right) peaks. (B) Bargraph showing genome wide distribution of Med1 bindings. (C) Med1 was used to identify stage specific super-enahncers by ROSE. (D) Venn diagram showing the number of naïve, primed and shared Med1 super-enahncers. (E) Venn diagram showing number of Med1 super-enhancers overlapped with stage-specific AT-dep MLL3/4, Brg1-H3K27ac, and Brg1-H3K27ac-MLL3/4+MLL3/4-H3K3K27ac enhancers. (F) Bargraph showing the percentage of AT-hook dependent differential genes (Pausing) that are within 50kb, 100kb, 250kb, 500kb, 750kb, and 1Mb distance from Med1 naïve and primed specific super-enhancers.

### AT-hook domain is required for cell lineage priming but not stem cell maintenance

Next, we examine whether the large-scale transcription changes observed in naïve cells alters stem cell maintenance and proliferation and the expression/localization of the core pluripotency TFs. We had shown previously two classes of Oct4, Sox2, Nanog and Tcf binding sites by ATAC-seq. One group of these TF binding sites is only bound in naïve cells which is lost when the AT-hook is deleted. The other group of TF binding sites are bound in both naïve and primed cells and are those expected to control the core pluripotency transcription circuit given naïve and primed cells are both pluripotent. In the dAT mutants, we observe no defects in stem cell proliferation, clonal expansion, alkaline phosphatase staining or colony morphology (Figures 7A-C and S8A). Deletion of the AT-hook did not alter the expression or nuclear localization of pluripotency TFs Oct4, Nanog, and Sox2 by immunofluorescence (Figure 7D). The AT-hook of BRG1 is therefore not required to maintain the core pluripotency circuitry and associated transcription factor binding. We also observe the AT-hook is not required for proper SMARCA4 expression, nuclear localization, or complex integrity in either naïve or primed cells (Figures S8B-D). The AT-hook of BRG1 appears to facilitate transcription in naïve cells that is distinct from that observed when BRG1 is absent (Kidder et al., 2009).

**Figure 7.**
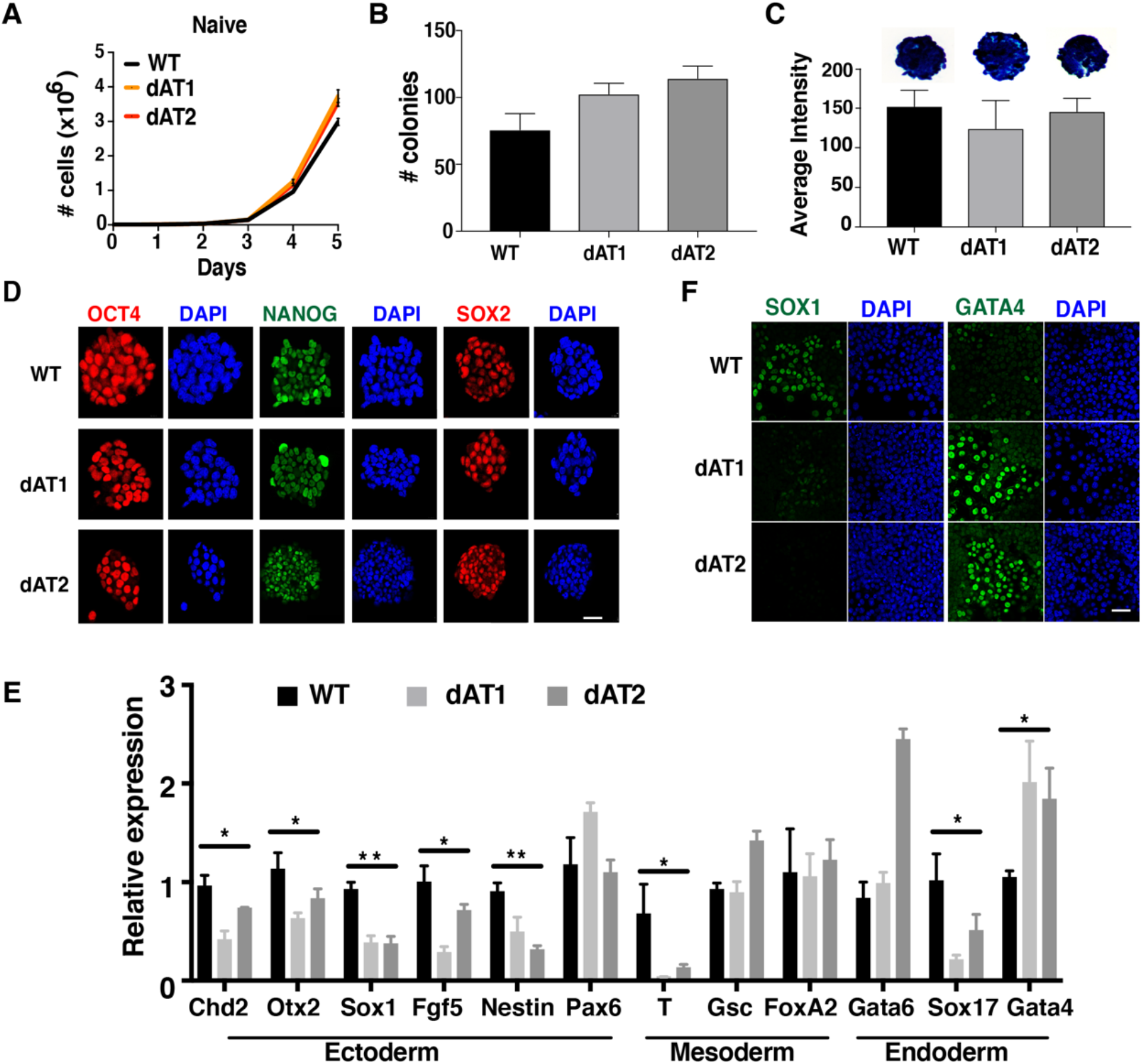
The AT-hook domain of SMARCA4 is required for cell lineage priming. **(A)** Growth curves of WT and AT-hook deletion mutant clones (dAT1, and dAT2) cultured in naïve condition are shown. (B-C) Bar graphs showing **the number of colonies formed** in a self-renewal assay (B), and average signal intensity of the colonies in alkaline phosphatase assay (C) of WT and dAT mESCs in naïve condition. **(D)** Immunofluorescence images showing expression and localization of the pluripotency markers OCT4, NANOG, and SOX2 in WT and AT-hook deleted mutant mESCs in naïve cells (scale bar, 20 micron). (E) Bar graph shows the expression of lineage-specific markers in WT and dAT mutants after culturing for seven days in the absence of LIF/2i. Gene expression analysis was done by quantitative reverse transcription PCR (qRT-PCR) and the values were normalized with *Gapdh*. Results are presented as means ± sd (n=3); *p<0.05; **p<0.001; (unpaired student’s t-test). (F) Immunofluorescence images showing expression of SOX1 (ectoderm lineage marker) and GATA4 (endoderm lineage marker) in WT and dAT mutants after seven days of LIF/2i withdrawal. Scale bar, 20 microns.

The role of BRG1 in cell lineage priming is followed by monitoring the expression of three distinct classes of lineage specific markers by qRT-PCR after culturing cells in the absence of LIF and two inhibitors for 7 days. Ectodermal markers Sox1 and Nestin, mesoderm marker Tbxt and the endoderm marker Sox17 were all down regulated in both dAT mutant clones compared to WT cells, consistent with defects in early differentiation (Figure 7E). The endoderm specific gene Gata4 was aberrantly highly expressed in both dAT mutants compared to wild type (Figure 7E). The intercellular levels of Sox1 and Gata4 in the two dAT mutant clones as shown by immunofluorescence were respectively lower and higher than in wild type cells and suggests in the dAT mutant a bias toward differentiating into endoderm when compared to wild type (Figure 7F).

## Discussion

We have discovered a novel mode of SWI/SNF recruitment to cell-type specific cis-regulatory regions in naïve and primed mouse embryonic stem cells that involves a putative RNA binding module within the catalytic subunit called the AT-hook. The BRG1 catalytic subunit of the esBAF complex fails to bind cell-type-specific enhancers in two distinct pluripotent states when the AT-hook is deleted that also causes a redistribution of BRG1 to pre-existing BRG1 target sites. TF mediated recruitment of SWI/SNF is distinct from that mediated by the AT-hook and is independent of the AT-hook. We find BRG1 and its AT-hook are responsible for an early step in enhancer activation that precedes or promotes the acquisition of active histone marks (HK27ac and H3K4me1) and binding of some TFs and MLL3/4 that bridges enhancers to promoters. The stabilization of BRG1 binding mediated by p300/CBP and MLL3/4 co-activators is contingent on the AT-hook of BRG1. The activation of the catalytic activity of p300/CBP by BRG1 is therefore also strictly dependent on its AT-hook. MLL3/4 recruitment to enhancers is mediated by BRG1 and its AT-hook but differs from that of p300/CBP and H3K27ac AT-hook dependency in two ways. First, MLL3/4 recruitment mediated by the AT-hook does not promote monomethylation of H3K4 by MLL3/4; whereas stable recruitment of BRG1 mediated by the AT-hook facilitates acetylation of H3K27. Our data coincide well with other’s data that show MLL3/4 promotes transcription independent of its monomethyation of H3K4 (Dorighi et al., 2017). Second, BRG1 appears to be catalytically promoting MLL3/4 recruitment due to the low residency of BRG1 at these sites rather than direct physical interactions and is independent of its AT-hook in contrast to that observed for H3K27ac. The loss of TF binding that occurs when the AT-hook is deleted without changes in BRG1 co-localization is another indication that the AT-hook regulates the chromatin remodeling activity of BRG1. The AT-hook positively regulating BRG1 remodeling activity for TF recruitment is consistent with a prior study showing the catalytic activity of BRG1 being crucial for TF binding using chemical inhibitors that target the ATPase activity of BRG1 (Iurlaro et al., 2021). There are other sites where MLL3/4 is transiently bound that depend on AT-hook of BRG1 for the catalytic activity of MLL3/4. Monomethylation of H3K4 at these low MLL/3/4 residency sites depend on the AT-hook of BRG1 and suggests BRG1 can increase MLL3/4 catalytic activity when it is more rapidly turning over on chromatin.

BRG1 and its AT-hook activation of enhancers extends to regulation of cell-type specific super-enhancers. The majority of naïve and primed specific super-enhancers defined by high levels of MED1 and a clustering of MED1 sites have multiple regions where H3K27ac alone, MLL3/4 alone or both MLL3/4 and H3K27ac require BRG1 and its AT-hook. These super-enhancers vary in their composition of these elements and on the average have as many as 9 of these elements per super-enhancer. We find the MED1 subunit of the Mediator complex preferentially associates with enhancers relative to promoters and suggest a preferred discrete spatial orientation of Mediator when bridging promoters and enhancers. We find the genes that are differentially regulated in either naïve or primed cells when the AT-hook is deleted are located in relatively close proximity (250 kb-1 Mb) to the super-enhancers that are also differentially regulated by loss of the AT-hook, consistent with their co-regulation. The BRG1 regulated super-enhancers appear to regulate genes important for early cell lineage priming as well as some features of the naïve pluripotent state. The co-dependency of BRG1 and H3K27ac (p300/CBP) for their localization in pluripotency and early cell lineage priming is consistent with prior data showing them working together in cardiomyocyte differentiation (Alexander et al., 2015), but is the first time the AT-hook has been shown to be involved along with the implication of RNA being a crucial factor. Like what we observe in pluripotency and early cell lineage priming, BRG1 and MLL4 have been shown to reciprocally depend on each other for their binding to enhancers during adipogenesis (Park et al., 2021). BRG1 as a master regulator of super-enhancers and its connection to RNA is intriguing given the recent evidence of RNA playing a central part in the formation of transcription related condensates (Henninger et al., 2021; Sharp et al., 2022). Additional studies are needed to confirm the RNAs bound to the AT-hook of BRG1, if they function in a trans-or cis-manner to regulate BRG1 recruitment and/or chromatin remodeling activity, and if loss of the AT-hook of BRG1 impacts condensate formation.

## STAR Methods

### Key Resource Table

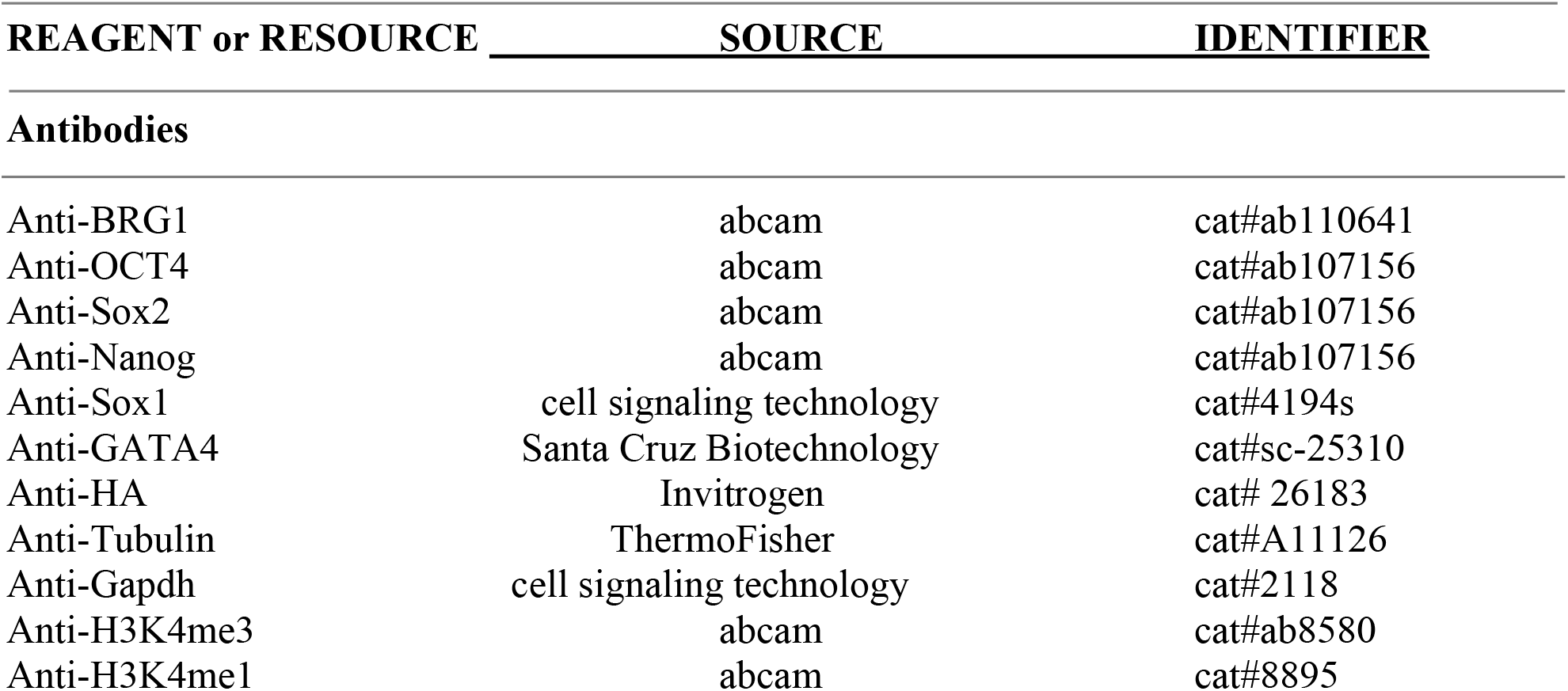
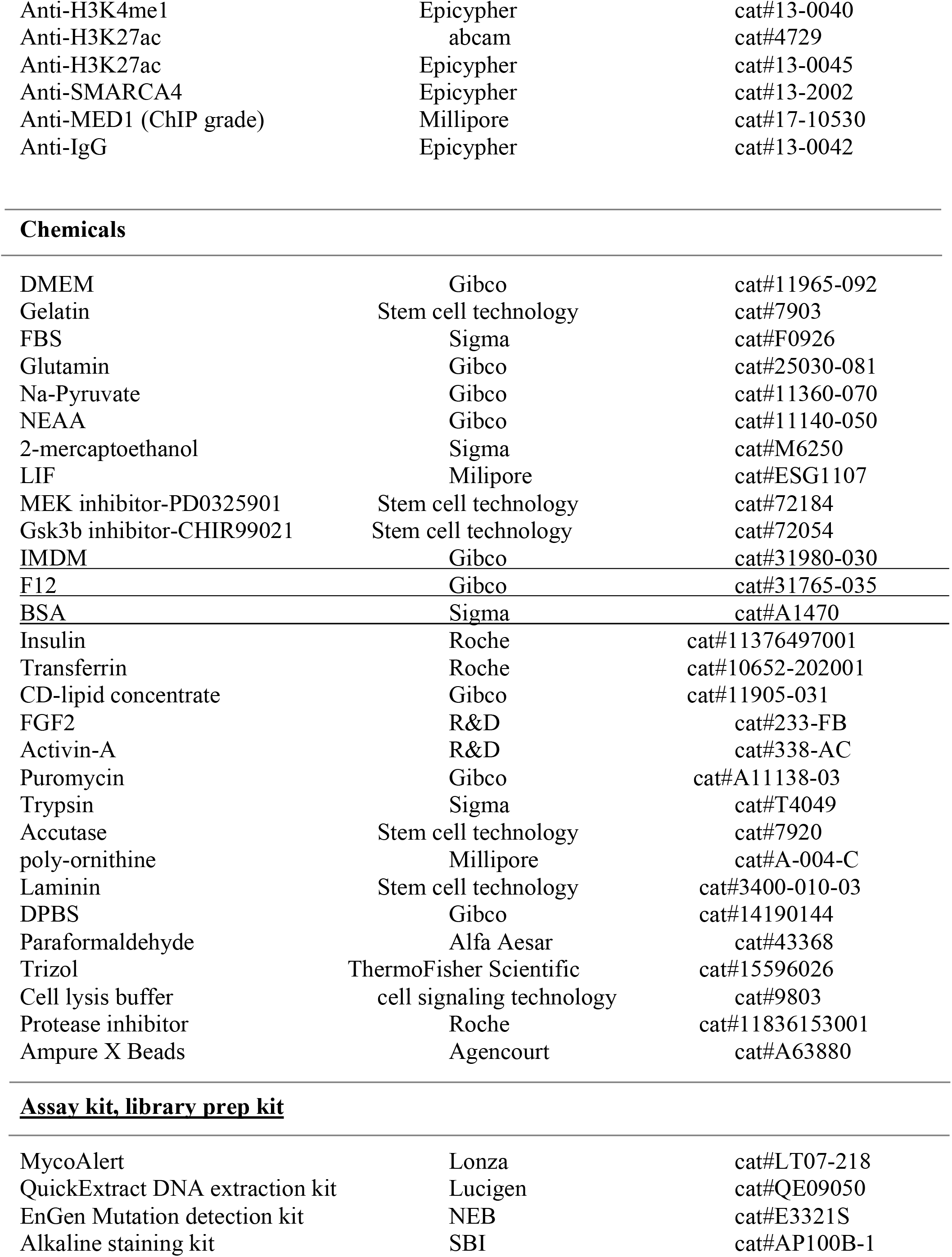
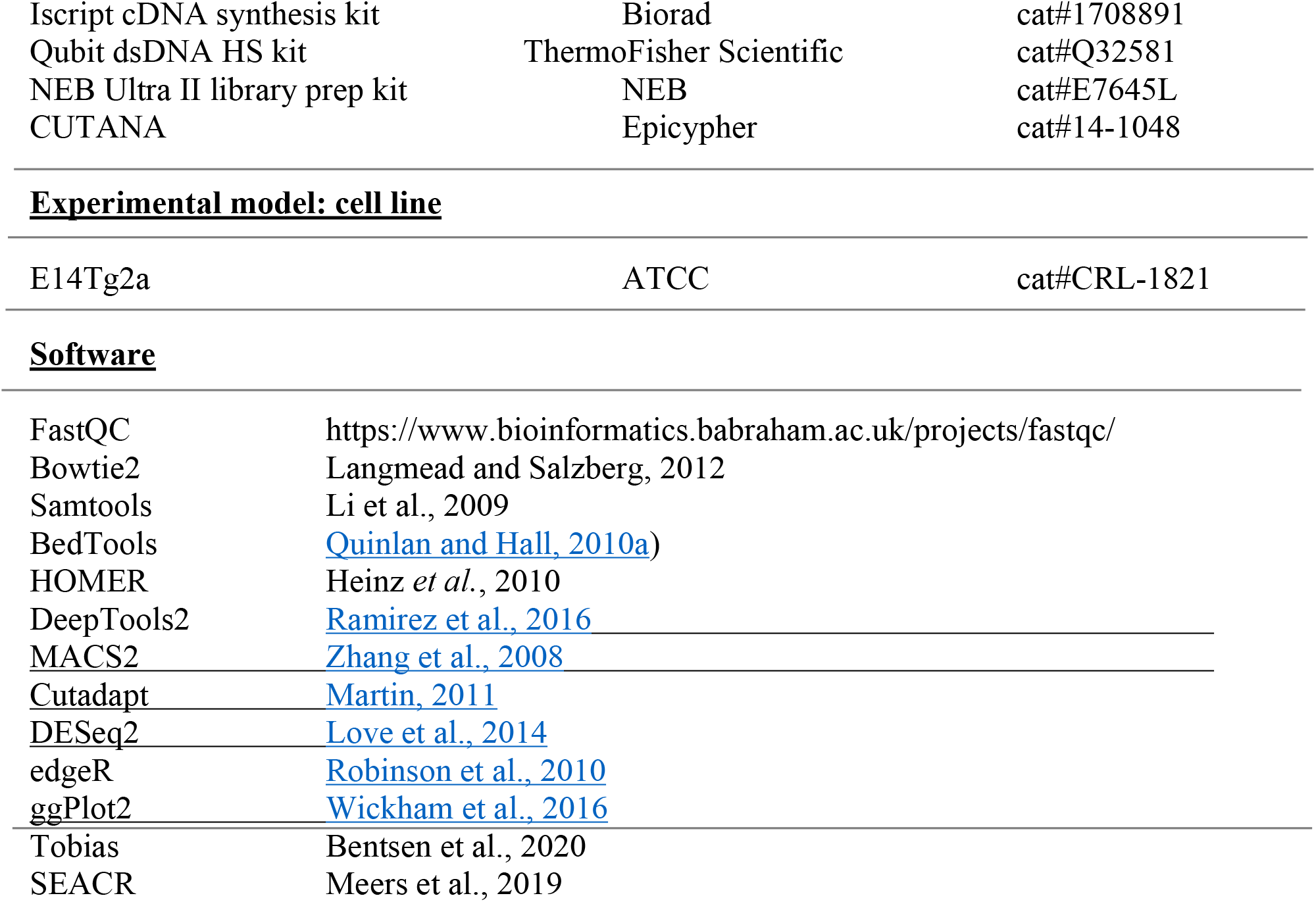

| REAGENT or RESOURCE | SOURCE | IDENTIFIER |
| --- | --- | --- |
| **Antibodies** |  |  |
| Anti-BRG1 | abcam | cat#ab110641 |
| Anti-OCT4 | abcam | cat#ab107156 |
| Anti-Sox2 | abcam | cat#ab107156 |
| Anti-Nanog | abcam | cat#ab107156 |
| Anti-Sox1 | cell signaling technology | cat#4194s |
| Anti-GATA4 | Santa Cruz Biotechnology | cat#sc-25310 |
| Anti-HA | Invitrogen | cat# 26183 |
| Anti-Tubulin | ThermoFisher | cat#A11126 |
| Anti-Gapdh | cell signaling technology | cat#2118 |
| Anti-H3K4me3 | abcam | cat#ab8580 |
| Anti-H3K4me1 | abcam | cat#8895 |
| Anti-H3K4me1 | Epicypher | cat#13-0040 |
| Anti-H3K27ac | abcam | cat#4729 |
| Anti-H3K27ac | Epicypher | cat#13-0045 |
| Anti-SMARCA4 | Epicypher | cat#13-2002 |
| Anti-MED1 (ChIP grade) | Millipore | cat#17-10530 |
| Anti-IgG | Epicypher | cat#13-0042 |
| Chemicals |  |  |
| DMEM | Gibco | cat#11965-092 |
| Gelatin | Stem cell technology | cat#7903 |
| FBS | Sigma | cat#F0926 |
| Glutamin | Gibco | cat#25030-081 |
| Na-Pyruvate | Gibco | cat#11360-070 |
| NEAA | Gibco | cat#11140-050 |
| 2-mercaptoethanol | Sigma | cat#M6250 |
| LIF | Millipore | cat#ESG1107 |
| MEK inhibitor-PD0325901 | Stem cell technology | cat#72184 |
| Gsk3b inhibitor-CHIR99021 | Stem cell technology | cat#72054 |
| IMDM | Gibco | cat#31980-030 |
| F12 | Gibco | cat#31765-035 |
| BSA | Sigma | cat#A1470 |
| Insulin | Roche | cat#11376497001 |
| Transferrin | Roche | cat#10652-202001 |
| CD-lipid concentrate | Gibco | cat#11905-031 |
| FGF2 | R&D | cat#233-FB |
| Activin-A | R&D | cat#338-AC |
| Puromycin | Gibco | cat#A11138-03 |
| Trypsin | Sigma | cat#T4049 |
| Accutase | Stem cell technology | cat#7920 |
| poly-ornithine | Millipore | cat#A-004-C |
| Laminin | Stem cell technology | cat#3400-010-03 |
| DPBS | Gibco | cat#14190144 |
| Paraformaldehyde | Alfa Aesar | cat#43368 |
| Trizol | ThermoFisher Scientific | cat#15596026 |
| Cell lysis buffer | cell signaling technology | cat#9803 |
| Protease inhibitor | Roche | cat#11836153001 |
| Ampure X Beads | Agencourt | cat#A63880 |
| Assay kit, library prep kit |  |  |
| MycoAlert | Lonza | cat#LT07-218 |
| QuickExtract DNA extraction kit | Lucigen | cat#QE09050 |
| EnGen Mutation detection kit | NEB | cat#E3321S |
| Alkaline staining kit | SBI | cat#AP100B-1 |
| Iscript cDNA synthesis kit | Biorad | cat#1708891 |
| Qubit dsDNA HS kit | ThermoFisher Scientific | cat#Q32581 |
| NEB Ultra II library prep kit | NEB | cat#E7645L |
| CUTANA | Epicypher | cat#14-1048 |
| **Experimental model: cell line** |  |  |
| E14Tg2a | ATCC | cat#CRL-1821 |

**Software**
|  |  |
| --- | --- |
| FastQC | https://www.bioinformatics.babraham.ac.uk/projects/fastqc/ |
| Bowtie2 | Langmead and Salzberg, 2012 |
| Samtools | Li et al., 2009 |
| BedTools | Quinlan and Hall, 2010a) |
| HOMER | Heinz *et al.*, 2010 |
| DeepTools2 | Ramirez et al., 2016 |
| MACS2 | Zhang et al., 2008 |
| Cutadapt | Martin, 2011 |
| DESeq2 | Love et al., 2014 |
| edgeR | Robinson et al., 2010 |
| ggPlot2 | Wickham et al., 2016 |
| Tobias | Bentsen et al., 2020 |
| SEACR | Meers et al., 2019 |

### Resource availability

Further information and requests for resources and reagents should be directed to and will be fulfilled by the lead contact, Blaine Bartholomew.

### Experimental Model and subject details

#### Cell culture

Mouse embryonic stem cells (E14Tg2a, ATCC) were maintained on 1% gelatin-coated plates in the ESGRO complete plus clonal grade medium (Millipore), as described(Cinghu et al., 2014; Oldfield et al., 2014). Embryonic stem cells (ESCs) were cultured on gelatin-coated plates in DMEM (Invitrogen) supplemented with 15% FBS, 1X-Gultamax (Gibco), Na-Pyruvate (Gibco), 10mM 2-mercaptoethanol, 0.1 mM nonessential amino acids (Gibco), 1U/ml of ESGRO mLIF (Millipore), and 2i inhibitors (MEK inhibitor PD0325901, Gsk3b inhibitor (CHIR99021 – Stem Cell technology) in naïve condition. In epiblast stem cells (EpiSCs) or primed condition, cells were cultured in chemically defined medium (IMDM and F12-Invitrogen) supplemented with 2%-BSA (Sigma), Insulin (Roche), Transferrin (Roche), CD-lipid concentrate (Gibco), FGF2 (R&D), and Activin-A (R&D) (Ref). For spontaneous differentiation (SD), cells were cultured in gelatin coated plate in previously described DMEM-FBS media without LIF and 2i-inhibitors. Cell lines were continuously monitored under microscope and confirmed to be free of mycoplasma contamination by using a MycoAlert mycoplasma detection kit (Lonza) and DAPI staining.

### Method Details

#### Generation of stable constructs

CRISPR mediated deletion of the AT-hook of SMARCA4/BRG1 and insertion of the HA-tag Guide RNA used for CRISPR-Cas9 editing were designed using the CRISPR Design Tool (http://crispr.mit.edu) to minimize off-target effects. G-blocks containing the guide RNA directed to the AT-hook domain of SMARCA4/BRG1 (exon 30) came from IDT and were PCR amplified, cloned into two Cas9 containing plasmids (pX330; Addgene#158973) using Zero Blunt TOPO Cloning Kit (Invitrogen) and sequence verified. After 72 hrs of transfection in mESCs using electroporation, positive colonies were selected based on puromycin (1mg/ml) selection. Plasmids were separately tested in trial transfection in E14 cells to determine the efficiency of guide RNA cleavage by extracting genomic DNA (QuickExtract DNA Extraction from Epicentre). The regions was amplified by PCR and editing events identified using EnGen Mutation Detection Kit (NEB). All guide RNA used in HA-tag insertion or AT-hook deletion had a cleavage efficiency of >60%. ESCs were seeded at low density to allow for selection of individual colonies. Colonies were individually expanded and split for future culture or genomic DNA isolation. Genomic DNA (100 ng) from these colonies was used to confirm the desired targeted deletion by PCR. The donor DNA for HA-tag insertion contained a BamHI cut site that was used for screening positive clones. We obtained several clones with homozygous and heterozygous insertion of the HA-tag. We used a single BfuAI cut site in the wild type AT-hook region to screen for positive AT-hook deletion clones and from 129 clones obtained 10 homozygous knockouts of the AT-hook in E14 cells.

#### Cell Proliferation, Self-renewal assay and Alkaline phosphatase staining

Growth assay was done with 1×10^4^ cells seeded in gelatin coated six-well plates and monitored for the next 5 days in their respective culture condition (i.e. naïve and primed). Cell counting was done every other day using a hemocytometer and viability checked using trypan blue at the time of counting. Self-renewal and alkaline-phosphatase staining assays were done with 200 cells seeded in gelatin coated 12-wells plate and maintained in naïve media condition. Colonies were counted and stained with alkaline phosphatase (ALP) after 5 days. ALP staining was done following manufacturer’s protocol (SBI-AP100B-1). Media was changed every other day.

#### Immunofluorescence Staining

Cells were seeded in poly-ornithine (EMD-Millipore) and laminin (Sigma) coated 8-chambered slides (ibidi; cat#80806) and maintained in naïve and differentiation media independently. Next, cells were fixed with 4%-paraformaldehyde for 15 min at room temperature (RT) and washed with 1XPBS and permeabilized with 0.5% Triton for 5 min at RT. Blocking was done for 30 min at RT with 5% bovine serum albumin (Sigma). Primary antibodies were used with recommended dilutions and incubated for overnight at 4°C. After primary antibody incubation, cells were washed with 1XPBS (Phosphate buffer solution) and 1XPBST (PBS+0.1%Tween20) and then followed the secondary antibody incubation. For immunostaining, antibodies including anti-Oct4 (Abcam; ab107156), anti-Sox2 (Abcam; ab107156), anti-Nanog (Abcam; ab107156), anti-Brg1/SMARCA4 (Abcam; ab110641), anti-Sox1 (CST; 4194S), anti-Gata4 (Scbt; sc-25310) were used. Images were captured using a LSM-880 confocal microscope (Zeiss) and processed with Zen-blue software.

#### RNA isolation and quantitative real time PCR (qRT-PCR)

Total RNA was isolated using Trizol (Invitrogen), following the manufacturer’s protocol. One mg of RNA was used to prepare cDNA and were synthesized using the iScript kit (Bio-rad) according to the manufacturer’s protocol. For each biological replicate, quantitative PCR reactions were performed in technical triplicates using the iTaq Universal Sybr Green Supermix (Bio-Rad) on the Bio-Rad CFX-96 Real-Time PCR System, and the data normalized to *Gapdh*.

#### Immunopurification of wild type and dAT mutant SMARCA4/BRG1 complexes

Purification of HA-tag protein was performed following a previously described protocol (Gatchalian et al., 2018). Briefly, cells (∼2-3 × 10^8^) were cultured in gelatin coated plates and maintained in naïve and primed media independently. The packed cell volume (PCV) was estimated after cell harvesting and gently resuspended in buffer with 10mM HEPES-KOH pH 7.9, 10mM KCl, 1.5mM MgCl_2_ 1mM DTT, 1mM PMSF, 1uM pepstatin, 10uM chymostatin. Next, the cell suspension was transferred to a Dounce homogenizer fitted with a B-type pestle and the cells lysed with 20 strokes followed by centrifugation for 5 min at 900 x g. After removing the supernatant, the nuclei pellet was resuspended in buffer containing 0.2mM EDTA, 20% Glycerol, 20mM HEPES-KOH pH 7.9, 420mM NaCl, 1.5mM MgCl_2_, 1mM DTT, 1mM PMSF, 1uM pepstatin, 10uM chymostatin, protease inhibitor cocktail and the pellet lysed by gentle Dounce homogenization (i.e., 10-20 strokes with type-B pestle). The tube was mounted on a vortex mixer and agitated very gently for 30 minutes to 1hr at 4°C degree and centrifuged for 15 minutes at 20,000 x g and the supernatant collected. Nuclear lysates were diluted with two-thirds of original volume of 20mM HEPES, pH 7.9, and 0.3% NP-40 to adjust to an appropriate NaCl concentration. Anti-HA agarose beads were used following the manufacturer’s protocol (Mashtalir et al., 2020) and the nuclear extract was incubated with HA-beads overnight at 4ºC with gentle end-over-end mixing or on a rocking platform. Next, beads were washed three times with wash buffer (50mM Tris pH 8.0, 150mM NaCl, 1mM EDTA, 10% Glycerol, and 0.5% Triton X-100) and one bed volume of HA-peptide was added to the beads and incubated at 4ºC for 4 hr prior to elution of the complex. Untagged cells were lysed using 1X lysis buffer (CST) following manufacturer’s protocol to prepare the whole cell extract. The efficiency of the immunoprecipitation was checked by Western blotting following standard protocols with the following primary antibodies anti-Brg1/SMARCA4 (Abcam; ab110641), anti-HA (Invitrogen; Cat #26183), anti-Tubulin (ThermoFisher; A11126), and anti-Gapdh (CST; 2118). The immunoblot were visualized using Super Signal Pico chemiluminescent reagent.

#### PRO-seq

PRO-seq was performed as previously described with minor modifications(Mahat et al., 2016). Briefly, nuclei were isolated using Dounce homogenizer (1 million cells per mL) and nuclear run-on was performed with all four biotin-NTPs. RNAs were extracted by Trizol LS (Ambion) and fragmentated by base hydrolysis. From the fragmentated RNAs, biotin RNAs were enriched by streptavidin beads. The biotin RNAs were enriched twice more in each 3’ and 5’ RNA adaptor ligation processes. Reverse transcription, PCR amplification, and library size selection were performed to obtain 140 ∼ 350bp, 5ng/uL libraries. These libraries were sequenced on an Illumina NextSeq 500 (75bp, single-end reads).

#### ChIP-seq

Chromatin immunoprecipitation (ChIP) was performed following previously described high-throughput ChIP protocol with some modifications (Blecher-Gonen et al., 2013). Cells were crosslinked in 1% formaldehyde for 10 min at room temperature, before quenching with 125 mM glycine for 5 min. After 2 washes with ice cold PBS, cells were incubated in lysis buffer (5 mM PIPES pH 8.0, 85 mM KCl, 0.5 % NP-40 supplemented with protease inhibitor) for 10 min, and nuclei were collected after centrifugation. The nuclear pellet was re-suspended in shearing buffer (12 mM Tris-HCl pH 7.5, 6 mM EDTA pH 8.0, 0.5 % SDS supplemented with protease inhibitor) and chromatin was fragmented using ME220 focused ultra-sonicator (Covaris) to obtain DNA fragments ranging 200-600 bp. The chromatin lysate was collected after centrifugation and incubated overnight at 4°C with SMARCA4 and respective histone antibodies conjugated with Dynabeads Protein G (Invitrogen). Next day, antibody-bound DNA were collected using Dynamag, washed extensively as described in the protocol, treated with RNase and Proteinase K, and reverse crosslinked overnight followed by DNA extraction using Ampure X beads (Beckman Coulter). Purified ChIP DNA was used for library construction using NEB Ultra II DNA library prep kit (New England Biolabs) and submitted for sequencing (75bp paired end and 50 bp single reads) on an Illumina HiSeq3000.

#### ATAC-seq

ATAC-seq was performed as previously described(Buenrostro et al., 2013). Briefly, 50,000 cells were washed with cold PBS, collected by centrifugation then resuspended in resuspension buffer (10 mM Tris-HCl, pH 7.4, 10 mM NaCl, 3 mM MgCl2). After collection, cells were lysed in lysis buffer (10 mM Tris-HCl, pH 7.4, 10 mM NaCl, 3 mM MgCl2, 0.1% NP-40) and collected before incubating in transposition mix containing Tn5 transposase (Illumina). Purified DNA was then ligated with adapters, amplified and size selected for sequencing. Library DNA was sequenced with paired end 50 bp reads.

#### CUT&RUN

CUT&RUN for MLL3/4, BRG1, H3K27ac and H3K4me1 in E14 WT and mutant ES cells was done using CUTANA ChIC/CUT&RUN kit (Epicypher #14-1048) following manufacturer’s protocol. Briefly, 5 × 10^5^ cells were captured with Concanavalin A, permeabilized using 0.01% digitonin and incubated with 0.5 μg antibody (anti-MLL3/4 serum [kind gift from Dr. Joanna Wysocka]/ anti-BRG1[#13-2002]/ anti-H3K27ac [#13-0045]/ anti-H3K4me1 [#13-0040; Epicypher]/ IgG [# 13-0042; Epicypher]) in 50 μL antibody buffer (20 mM HEPES at pH 7.5, 150 mM NaCl, 0.5 mM Spermidine, 1x protease inhibitor cocktails [EDTA-free tablet; Roche], 2 mM EDTA, 0.01% digitonin) for overnight. After removing unbound antibody, pAG-MNase (20X) in 50 μL cell permeabilization buffer was added to the cells and incubated for 10 min at RT. Then CaCl_2_ (2 mM) was added to activate MNase and incubated at 4 °C for 2 hr. The reaction was stopped using 33 μL of 2X STOP buffer (340 mM NaCl, 20 mM EDTA, 4 mM EGTA, 50 μg/mL RNase A, 50 μg/mL glycogen) and *E. coli* spike-in DNA (0.5ng) added as a control. The DNA from the released chromatin in the supernatant was purified and then quantified using Qubit dsDNA HS assay kit (Agilent Technologies). CUT&RUN libraries were constructed using NEBNext Ultra II DNA library preparation kit as described previously(Skene et al., 2018) with a few modifications. Briefly, end repair and dA-tailing were conducted on 6 ng of CUT&RUN eluted DNA for 30 min at 20°C followed by 30 min at 65°C. After adaptor ligation for 30 min at 20°C, the DNA fragments were purified by 1X vol of AMPure XP beads (Beckman Coulter) followed by 10-12 cycles of PCR amplification with Next Ultra II Q5 master mix. The PCR products were purified with 1X vol of AMPure XP beads. After quantitative and qualitative analysis, libraries with different indexes were pooled and sequenced on Illumina HiSeq3000 platform with paired-end 75-bp reads.

#### FASP Methods – Orbitrap Exploris DIA

Protein samples were reduced, alkylated, and digested using filter-aided sample preparation with sequencing grade modified porcine trypsin (Promega). Tryptic peptides were then separated by reverse phase XSelect CSH C18 2.5 um resin (Waters) on an in-line 150 × 0.075 mm column using an UltiMate 3000 RSLCnano system (Thermo). Peptides were eluted using a 60 min gradient from 98:2 to 65:35 buffer A:B ratio (Buffer A = 0.1% formic acid, 0.5% acetonitrile; Buffer B = 0.1% formic acid, 99.9% acetonitrile). Eluted peptides were ionized by electrospray (2.2 kV) followed by mass spectrometric analysis on an Orbitrap Exploris 480 mass spectrometer (Thermo). To assemble a chromatogram library, six gas-phase fractions were acquired on the Orbitrap Exploris with 4 m/z DIA spectra (4 m/z precursor isolation windows at 30,000 resolution, normalized AGC target 100%, maximum inject time 66 ms) using a staggered window pattern from narrow mass ranges using optimized window placements. Precursor spectra were acquired after each DIA duty cycle, spanning the m/z range of the gas-phase fraction (i.e., 496-602 m/z, 60,000 resolution, normalized AGC target 100%, maximum injection time 50 ms). For wide-window acquisitions, the Orbitrap Exploris was configured to acquire a precursor scan (385-1015 m/z, 60,000 resolution, normalized AGC target 100%, maximum injection time 50 ms) followed by 50x 12 m/z DIA spectra (12 m/z precursor isolation windows at 15,000 resolution, normalized AGC target 100%, maximum injection time 33 ms) using a staggered window pattern with optimized window placements. Precursor spectra were acquired after each DIA duty cycle.

### Data-analysis

#### PRO-seq analysis

Adaptor trimming and low-quality reads were removed by Cutadapt(Martin, 2011). The filtered reads were aligned on mm10 genome by bowtie2(Langmead and Salzberg, 2012) with “--very-sensitive” option to discard reads mapped to more than one region. The mapped reads were compressed as binary form using samtools(Li et al., 2009) and rRNAs were removed from those reads by bedtools intersect(Quinlan and Hall, 2010a). The 3’ end of the filtered reads were captured in a strand-specific manner using bedtools genomecov. These files were used to calculate raw readcount of annotated genes by bedtools map. Similarity between biological replicates and two AT hook deletion clones, and difference between samples were confirmed by pearson correlation with clustering and PCA analysis using R package DESeq2(Love et al., 2014). After the validation of replicates, all replicates were merged and normalized to meet final 30 million reads. The normalized reads were used to obtain differential genes through log2 fold-change calculation, create MA plots using R package ggplot2(Wickham et al., 2016) and perform meta-plot analysis using deepTools2(Ramirez et al., 2016) with gaussian smoothing by R package Smoother(Spiess et al., 2015). Statistically significant gene ontology (GO) terms for each differential genes were annotated by GO enrichment analysis (http://geneontology.org/) (Ashburner et al., 2000; Consortium, 2020; Mi et al., 2018).

#### Gene list identification

Annotation of TSSs and TESs for 55,487 genes are downloaded from Gencode vM22 and active TSSs for mESC identified through START-seq are provided by Dr. Karen Adelman’s lab (Henriques et al., 2018). If active TSS are located within ±1kb region from the TSS of the 55,487 genes, then the TSS of this gene is replaced by the position of this active TSS. Genes with length (TSS-TES) < 2kb and without PRO-seq signal at either promoter proximal pausing (TSS-100bp + TSS+300bp) or early elongating regions (TSS+300bp ∼ TSS+2kbs) were removed to avoid genes with unclear transcripts. Thus, 15,265 genes with the length of transcription (TSS-TTS) > 2kb with valid expression remained for further study.

#### ATAC-seq, ChIP-seq, and CUT&RUN

Data from two biological replicates were first compared (R^2^ > 0.9), and then merged into a single read file for each time point. ATAC-seq peaks were then called using MACS2 (Zhang et al., 2008) with the following parameters: -q 0.01–nomodel–shift 75 –extsize 150. To get a union set of peaks from all samples (WT and dAT mutants), MACS2 peaks from each condition were merged using mergePeaks module from HOMER (Heinz et al., 2010) (default parameters). For identifying differentially accessible regions, the union set of peaks was annotated by Homer and then divided into promoter (−1 kb to +1 kb), and intronic-intergenic regions. Read counts for all peaks in the union set were obtained using the featureCount module of Subread package (Liao et al., 2014) and differential anlysis was done using edgeR (Robinson et al., 2010). Heatmaps were created using deeptools (Ramirez et al., 2016) plotHeatmap function. Data from two biological replicates were first compared (R^2^ > 0.9), and then merged into a single read file for each time point. ATAC-seq peaks were then called using MACS2 (Zhang et al., 2008) with the following parameters: -q 0.01– nomodel–shift 75 –extsize 150. To get a union set of peaks from all samples (WT and dAT mutants), MACS2 peaks from each condition were merged using mergePeaks module from HOMER (Heinz et al., 2010) (default parameters). For identifying differentially accessible regions, the union set of peaks was annotated by Homer and then divided into promoter (−1 kb to +1 kb), and intronic-intergenic regions. Read counts for all peaks in the union set were obtained using the featureCount module of Subread package (Liao et al., 2014) and differential analysis was done using edgeR (Robinson et al., 2010). Heatmaps were created using deeptools (default parameters). For identifying differentially accessible regions, the union set of peaks was annotated by Homer and then divided into promoter (−1 kb to +1 kb), and intronic-intergenic regions. Read counts for all peaks in the union set were obtained using the featureCount module of Subread package (Liao et al., 2014) and differential analysis was done using edgeR (Robinson et al., 2010). Heatmaps were created using deeptools (Ramirez et al., 2016) plotHeatmap function.

To determine the known motif enrichment findMotifsGenome module of the HOMER package was used. For motif heatmap analysis, differential ATAC-seq peaks between WT and dAT mutants were used to identify the known motifs using ‘findMotifsGenome.pl’ from Homer. Then, p-value of identified motifs were transformed into Z-score and plotted as a heatmaps using the R ggplot package. For ATAC footprint analysis, normalized ATAC files were corrected using ‘ATACorrect’ module from TOBIAS (Bentsen et al., 2020). Next, the average ATAC-signals were calculated (around +/-100bp of the center of the motif enriched peaks) and plotted using plotProfile from deeptools.

For ChIP-seq analysis, Paired-end 75 bp reads were aligned to mm10 using Bowtie 2 (Langmead and Salzberg, 2012) alignment tool using *--very-sensitive-local* preset parameter. Data from two biological replicates were first compared to check for concordance (R^2^ > 0.9), and then merged into a single read file for each cell type for further downstream analysis. The high confidence peak sets were selected from biological replicates using the intersectBed function from BEDTools (Quinlan and Hall, 2010b) with default parameters. For histone and Brg1 ChIP, peaks were called using MACS2 (Zhang et al., 2008) with the following parameters: --broad -q 0.05 –nomodel - extsize 500. To compare the signals between IP and input, ‘bdgcmp’ from MACS2 option was used. Peaks were annotated using the annotatePeaks module of HOMER package (Heinz et al., 2010). All the heatmaps were drawn using ‘plotHeatmap’ from deeptools (Ramirez et al., 2016).

All the Cut-Run data were sequenced in Paired-end 75 bp length and reads were aligned to mm10 using Bowtie2 (Langmead and Salzberg, 2012) alignment tool using *--very-sensitive-local* preset parameter. Each sample was normalized using 1million reads from *E. coli* spikein control. Data from two biological replicates were first compared to check for concordance (R2 > 0.9), and then merged into a single read file for each cell type for further downstream analysis. Cut-Run peaks were then called using SEACR (Meers et al., 2019) with ‘stringent’ ‘non’ ‘EFDR 0.01’ options in each sample. Peaks were annotated using the ‘annotatePeaks’ module of HOMER package (Heinz et al., 2010).

### Super-enhancer analysis

SEs were identified using ROSE (https://bitbucket.org/young_computation/rose) algorithm with some modifications. Med1 Cut&RUN peaks were stiched together computationally if they were within 12500bp of each other, though peaks having a Refseq promoter (+/-2.5kb) were excluded from stitching. These stitched enhancers were ranked by their Med1 signals. Super-enhancers were defined geometrically as those enhancers above the point at which the line y=x is tangent to the curve (Cut-off value for MED1 signal).

### Mass spectrometry

Following data acquisition, the data was searched using an empirically corrected library and a quantitative analysis was performed to obtain a comprehensive proteomic profile. Proteins were identified and quantified using EncyclopeDIA and visualized with Scaffold DIA using 1% false discovery thresholds at both the protein and peptide level (Searle et al., 2018). The UniProt database for Mus musculus was used for the database search. Protein exclusive intensity values were assessed for quality using ProteiNorm, a user-friendly tool for a systematic evaluation of normalization methods, imputation of missing values and comparisons of different differential abundance methods (Graw et al., 2020). Popular normalization methods were evaluated including log2 normalization (Log2), median normalization (Median), mean normalization (Mean), variance stabilizing normalization (Ritchie et al., 2015), (Chawade et al., 2014), quantile normalization (Quantile), Cyclic loess normalization (Cyclic Loess), global robust linear regression normalization (RLR), and global intensity normalization (Global Intensity). The individual performance of each method was evaluated by comparing of the following metrices: total intensity, Pooled intragroup Coefficient of Variation (PCV), Pooled intragroup Median Absolute Deviation (PMAD), Pooled intragroup estimate of variance (PEV), intragroup correlation, sample correlation heatmap (Pearson), and log2-ratio distributions. The data was normalized using Cyclic Loess and statistical analysis was performed using Linear Models for Microarray Data (limma) with empirical Bayes (eBayes) smoothing to the standard errors (Ritchie et al., 2015). Proteins with an FDR adjusted p-value < 0.05 and a fold change > 2 were considered to be significant.

## Acknowledgements

NGS sequencing was performed at the Science Park NGS Core, with support from CPRIT Core Facility Support Grant RP120348. Mass spectrometry was done at the University of Arkansas Medical Center Proteomics Core with support from NIH grant R24GM137786. Immunofluorescence was done at the Science Park imaging core, with support from FCCIC Core and FCCIC CPRIT grant RP170628. This work was supported by NIH grant R01GM108908. DS received support from the Center of Cancer Epigenetics at MD Anderson Cancer Center and SA from the Cockrell Foundation.

## Author Contributions

K.F. prepared the PRO-seq, ATAC-seq and BRG1 ChIP-seq samples. The H3K4me1, H3K4me3 and H3K27ac samples were prepared by K.F., D.S and A.J. Cell imaging, qRT-PCR and SMARCA4/BRG1 complex purification experiments were done by D.S. J.L performed the bioinformatic analysis of PRO-seq and D.S. did the bioinformatic analysis for ChIP-seq, CUT&RUN, and ATAC-seq. S.A. prepared the CUT&RUN samples and Y.C.L made and characterized all of the mESC clones with HA tag and knock-in of the AT-hook deletion using CRISPR-Cas9. Y.L. did all the initial processing of NGS data. B.L. assisted and supervised the bioinformatic analysis and B.B. supervised this work. B.B., D.S., J.L., B.L. and S.A. assisted in the manuscript preparation.

