## Supplemental figures for "An RNA binding module of SWI/SNF is required for activation of cell-type specific enhancers and super-enhancers in early development"

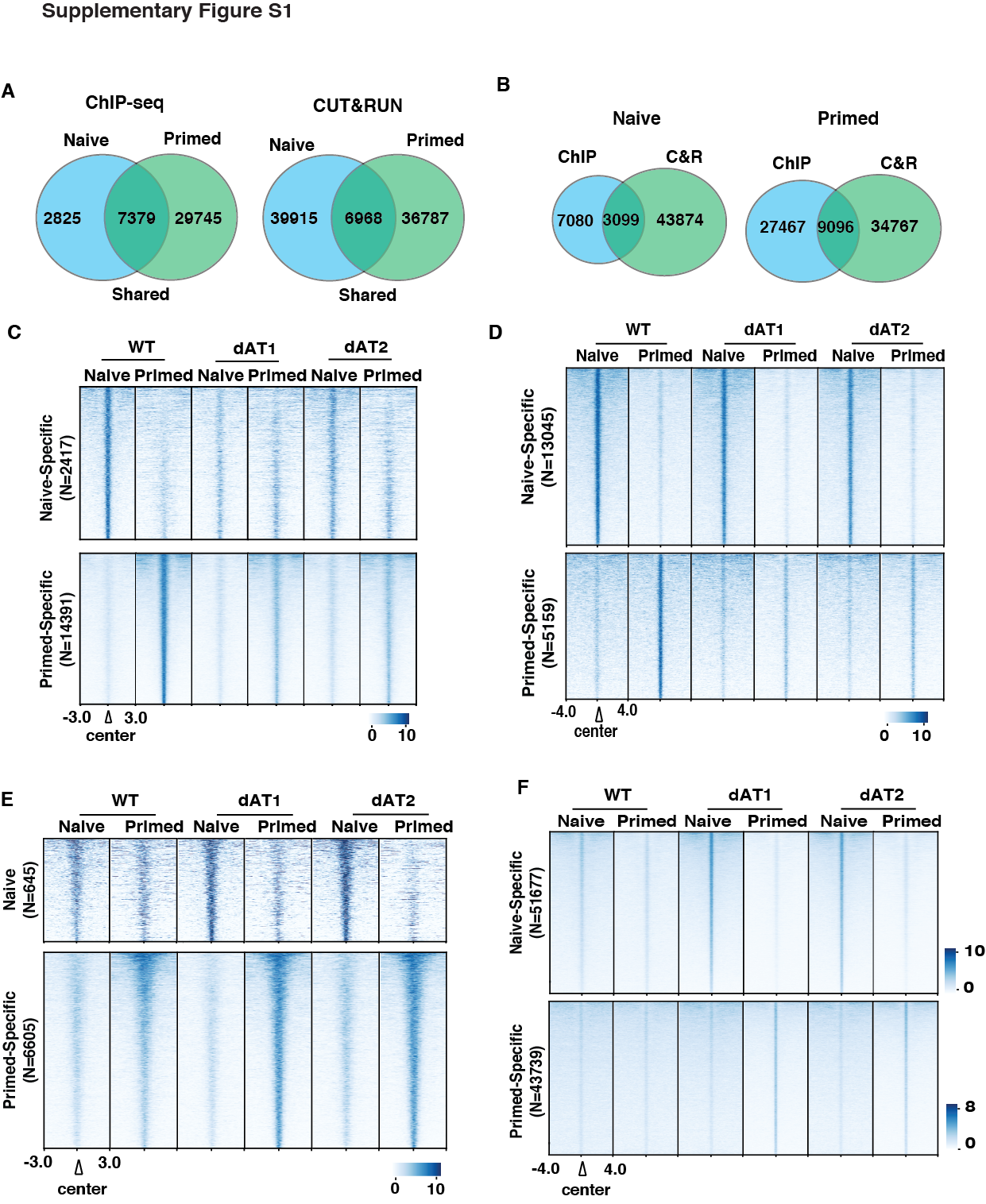

**Supplemental Figure S1. Redistribution of SMARCA4/BRG1 upon AT-hook deletion.**

(A) Venn diagrams show the number of BRG1 peaks detected in naïve and primed cells using ChIP-seq (left) and CUT&RUN (right). (B) The extent to which BRG1 peaks detected by ChIP-seq (ChIP) overlap with those of CUT&RUN (C&R) are shown. (C-D) BRG1 sites moderately dependent on the AT-hook for their localization are shown in these heatmaps using ChIP-seq (C) and CUT&RUN (D). (E-F) Some pre-existing BRG1 binding sites have increased levels BRG1 when the AT-hook is deleted as detected by ChIP-seq (E) and CUT&RUN (F) in naïve (upper panel) and primed (lower panel) cells.

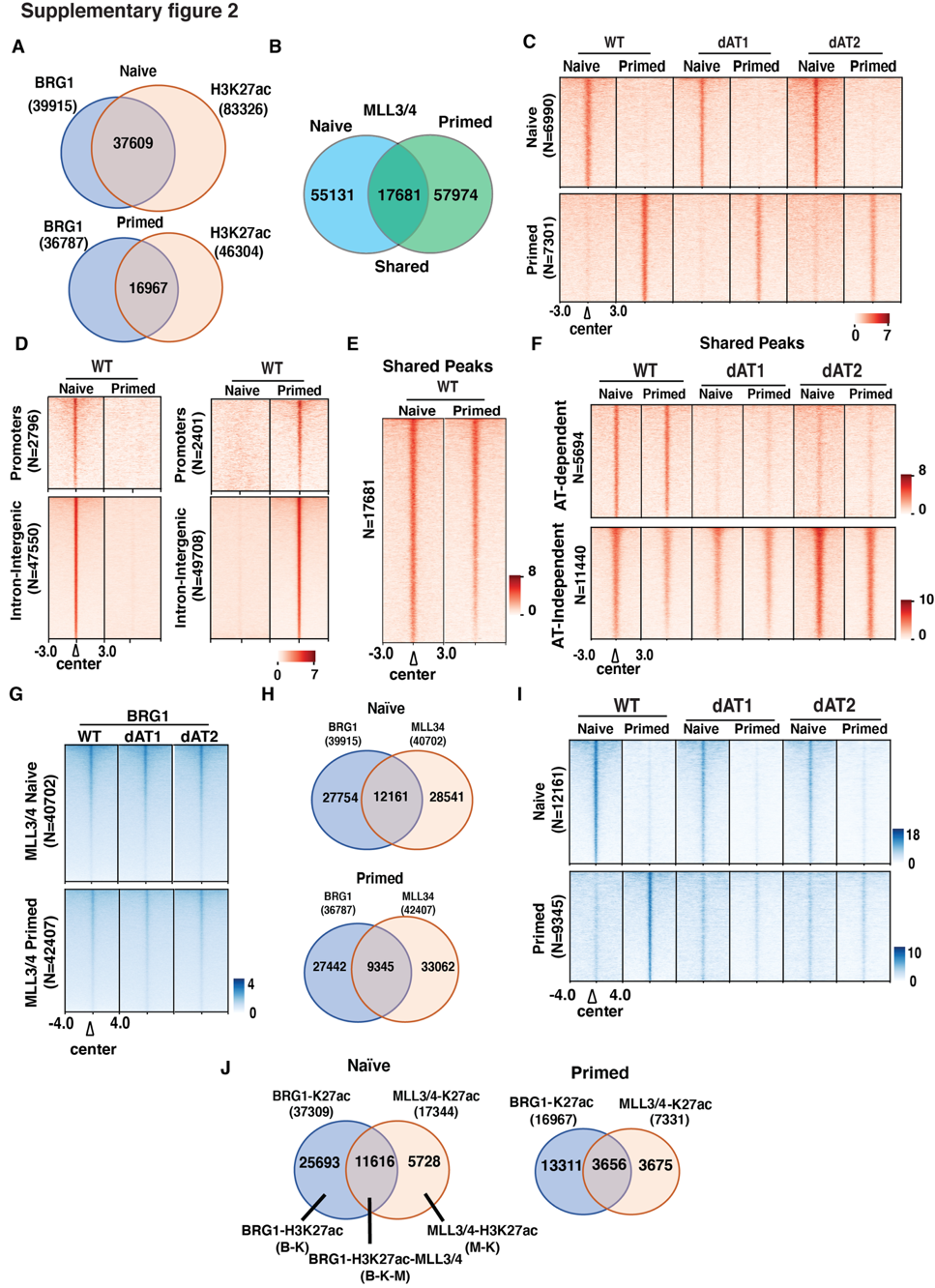

**Supplemental Figure S2. The role of the AT-hook of BRG1 in the interplay between BRG1, H3K27ac and MLL3/4**

(A) Venn diagrams the extent of overlap in the localization of BRG1 and H3K27ac in naïve and primed. (B) Venn diagram showing the MLL3/4 peaks detected in naïve and primed and the extent to which they are same in both cell types. (C) Heatmap show those MLL3/4 peaks that are not dependent on the AT-hook of BRG1 in naïve or primed. (D) The heat map shows the localization and distribution of MLL3/4 at promoter and cis-regulatory regions. (E-F) The MLL3/4 peaks that are shared between naïve and primed are shown in (E) and whether they are AT-dependent in (F). (G) BRG1 heatmap is shown for those regions where MLL3/4 are recruited to cis-regulatory regions in a naïve and primed-specific manner. (H) Shown is the extent of overlap between BRG1 and MLL3/4 peaks detected by CUT&RUN in naïve and primed. (I) BRG1 heatmap is shown for those regions where BRG1 and MLL3/4 co-localize as seen in (H). (J)The Venn diagram shows the extent of overlap between sites where BRG1 and H3K27ac colocalize versus those regions where MLL3/4 and H3K27ac co-localize and correlates with Figure 2C-D.

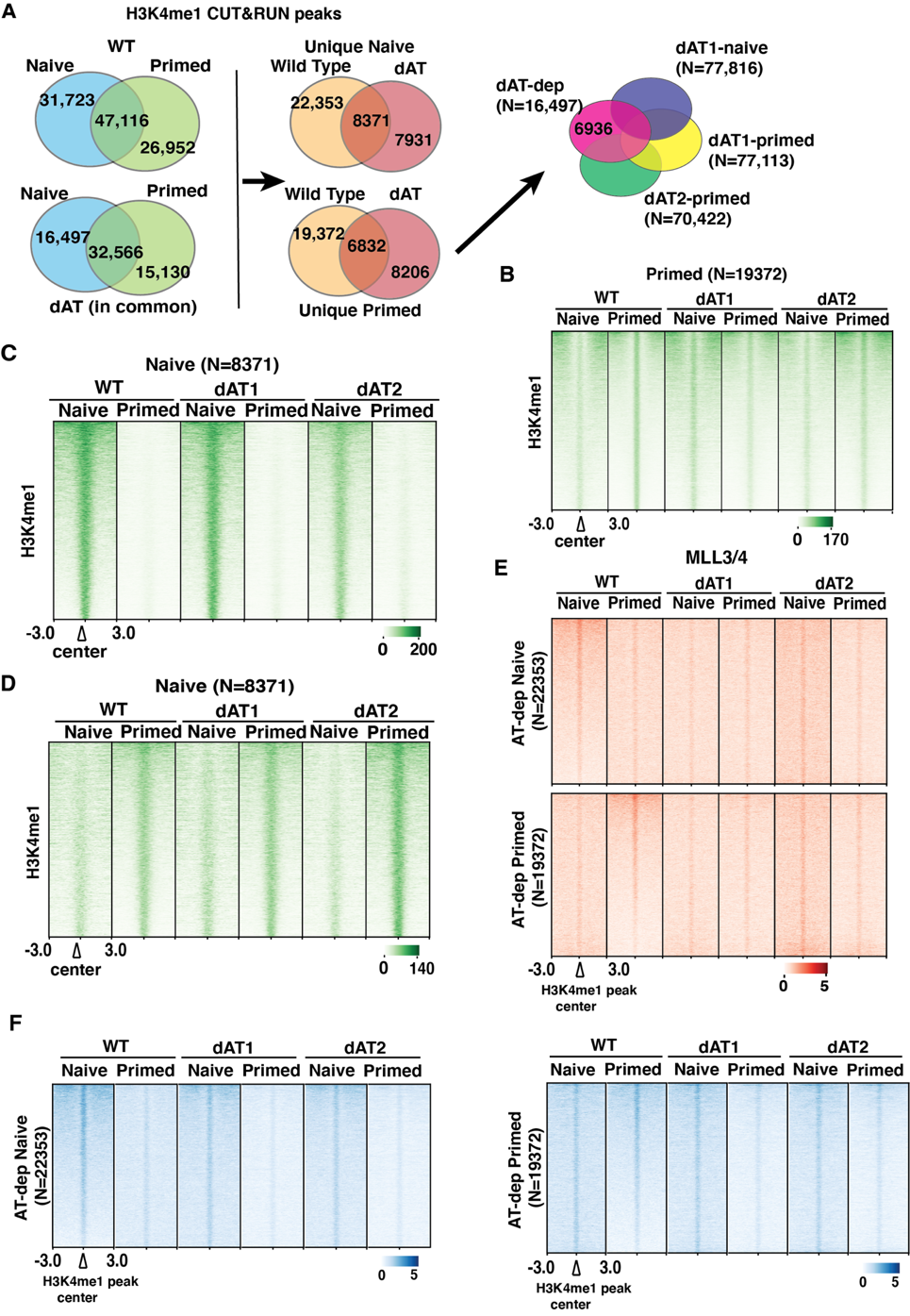

**Supplemental Figure S3. BRG1 and its AT-hook domain regulates H3K4me1 deposition.**

**(**A**)** Venn diagram showing naïve, primed, and shared H3K4me1 CUT&RUN peaks in WT (top left) and dAT mutants (bottom left). The Venn diagram shows H3K4me1 unique and shared peaks in WT and dAT mutants in naïve (top-middle) and primed (bottom-middle) states and unique AT-hook dependent H3K4me1 peaks in naïve (top right) and primed conditions (bottom right). (B) Heat map showing the primed-specific H3K4me1 peaks that are initially identified to be AT-hook dependent. (C-D) Heatmaps showing the H3K4me1 peaks that are not changed when the AT-hook is deleted in naïve (C) and primed (D) cells. (E-F) The profile of MLL3/4 (E) and BRG1 (F) at H3K4me1 peaks that are AT-hook dependent in naïve (upper panel) and primed (lower panel) cells. Signals are sorted based on WT in each group; N represents the total number of H3K4me1 intronic-intergenic peaks.

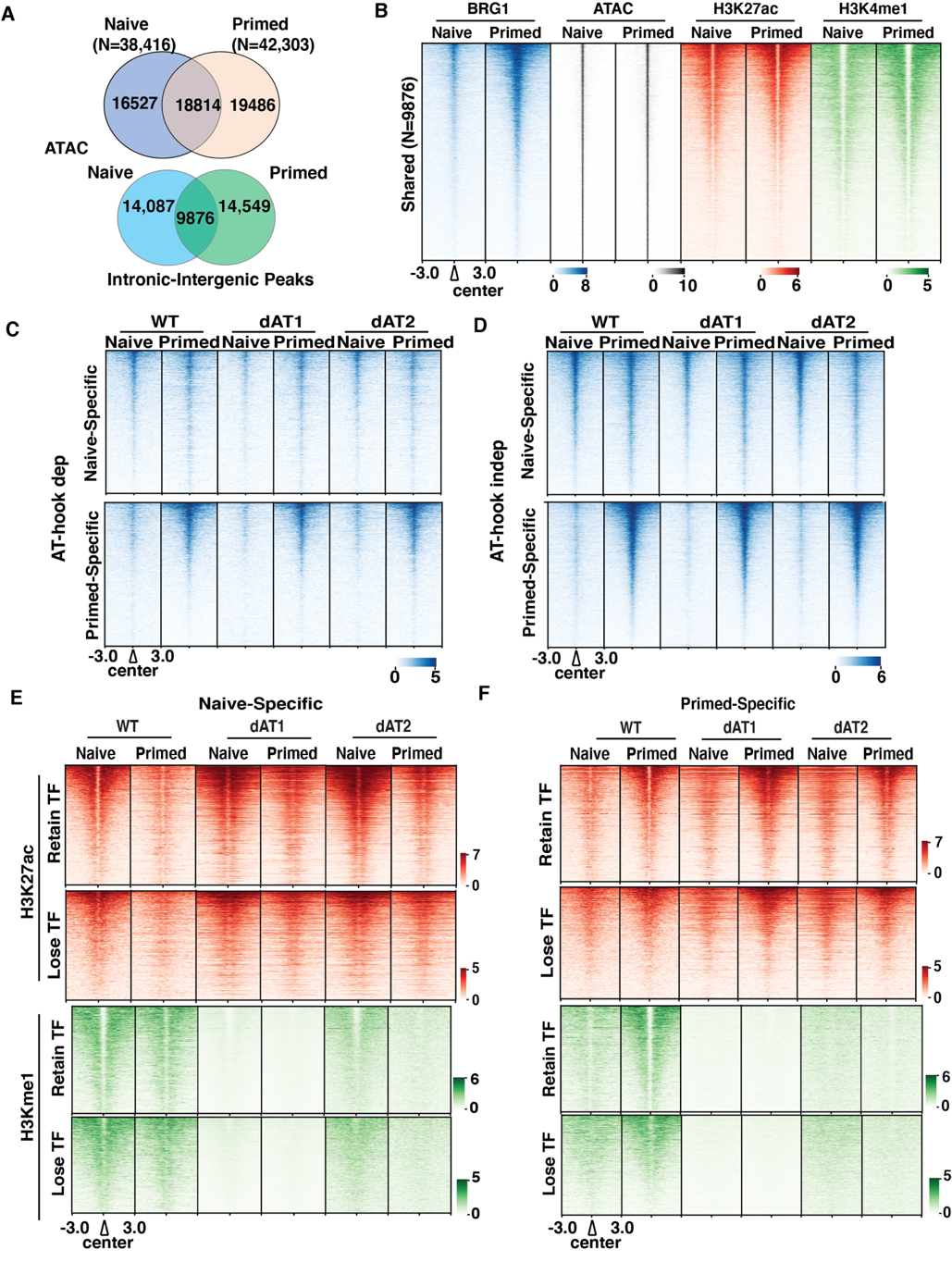

**Supplemental Figure S4. Deletion of AT-hook does not affect BRG1 localization at cis-regions. (**A**)** Venn diagrams show cell-type specific and shared ATAC-seq peaks (upper panel) and those only in intergenic and intronic regions (lower panel) in WT and dAT mutants in naïve and primed cells. (B) BRG1 localization (blue), ATAC-signal (grey), and active enhancer histone marks (H3K27ac [red], and H3K4me1 [green]) at shared intronic-intergenic ATAC-seq peaks (TF-dependent). Signals are sorted based on high to low. N represents the total number of ATAC-seq peaks. (C-D**)** Heatmap shows BRG1 localization at naïve (upper panel) and primed (lower panel) cis-regions where the ATAC signal changes when the AT-hook is deleted (C) or is not altered (D). (E-F) Heatmap shows ChIP-seq signals for H3K27ac (red, upper panel) and H3K4me1 (green, lower panel) at naïve (E) and primed (F) specific enhancers. The regions are divided into those where the ATAC accessibility is either AT-hook dependent (retain TF) or independent (loss of TF - bottom). ChIP signals are sorted based on WT in each group and based on naïve and primed WT in each group, respectively.

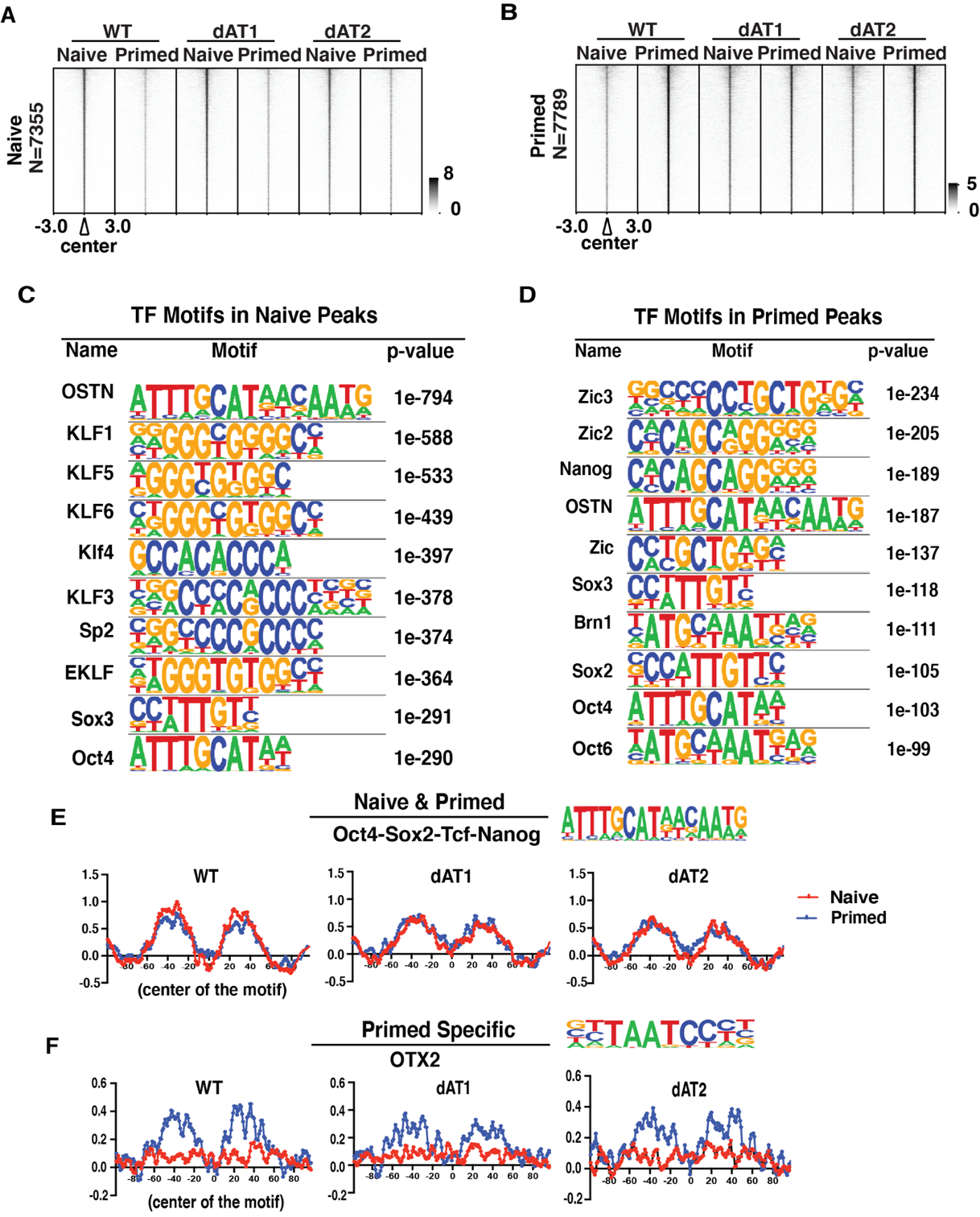

**Supplemental Figure S5. Pluripotency and epiblast-related transcription factors enriched in ATAC-seq peaks.**

**(A-B) Heatmap shows the ATAC peaks whose accessibility are not changed by the AT-hook** **like that shown in Figure 4D. (**C-D) Table showing the top 10 transcription factors motif enriched in naïve (C) and primed (D) ATAC-seq intronic-intergenic peaks using HOMER. (E-F) DNA footprinting of pluripotency OSTN TF (E) and EpiSC specific OTX2 TF (-F) binding sites that are unaltered in the dAT mutants.

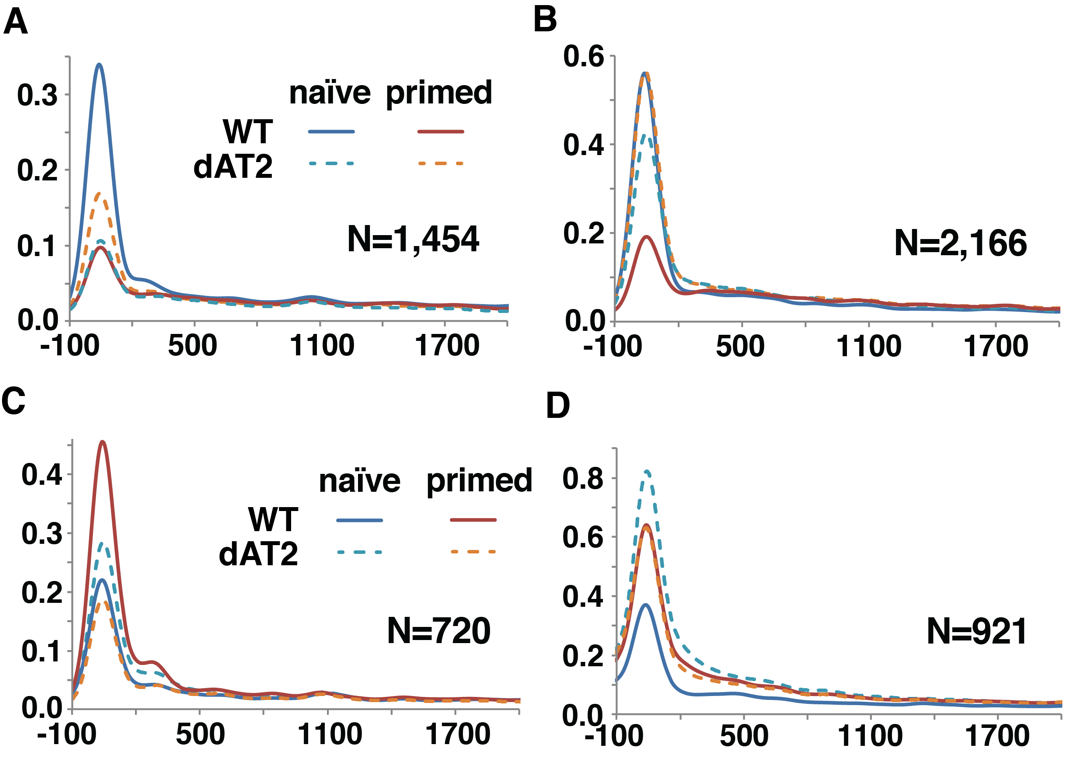

**Supplemental Figure S6. Transcriptional defects due to the loss of the AT-hook.** (A-B) Meta-analysis of PRO-seq signals (upstream TSS -100 bp to +300 bp downstream of TSS) for paused genes up-regulated in WT (A) and dAT2 mutant (B) in naïve condition. (C-D) Meta-analysis of PRO-seq signals (upstream TSS -100 bp to +300 bp downstream of TSS) for paused genes up-regulated in WT (A) and dAT2 mutant (B) in Primed condition.

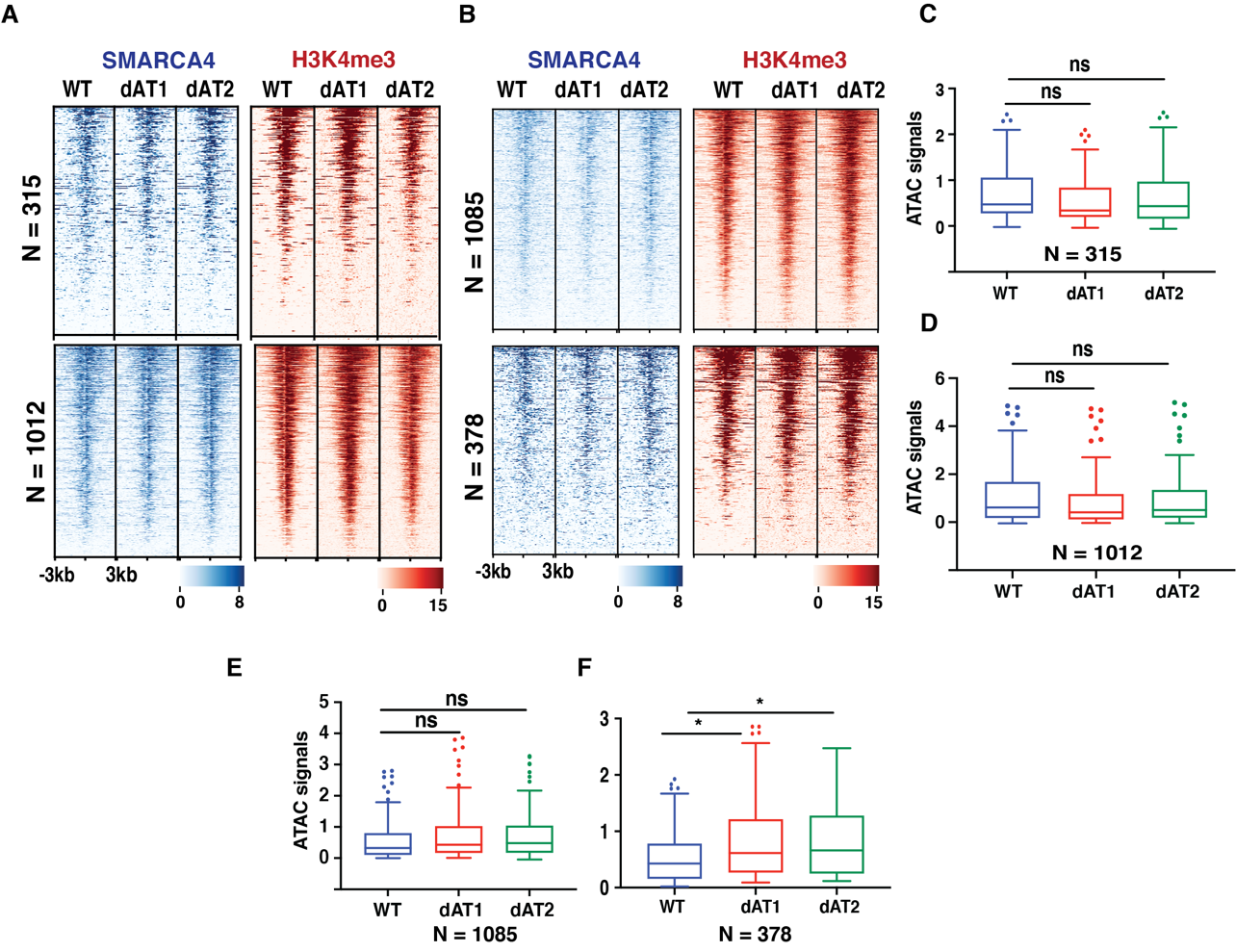
**Supplemental Figure S7. Deletion of AT-hook does not affect SMARCA4 localization and accessibility at promoter regions****.** (A-B**)** Heatmaps show localization of SMARCA4 (blue) and active histone mark H3K4m3 (red) in WT and AT-hook deletion clones at the promoter regions of differential genes that are being shared between dAT1 and dAT2 clones ChIP-signals are sorted based on WT SMARCA4. (C-F**)** Box plots showing average ATAC-seq signals in WT and dAT mutants at the promoter regions of genes. N represents the number of genes in each group.

**Supplementary Figure S8**

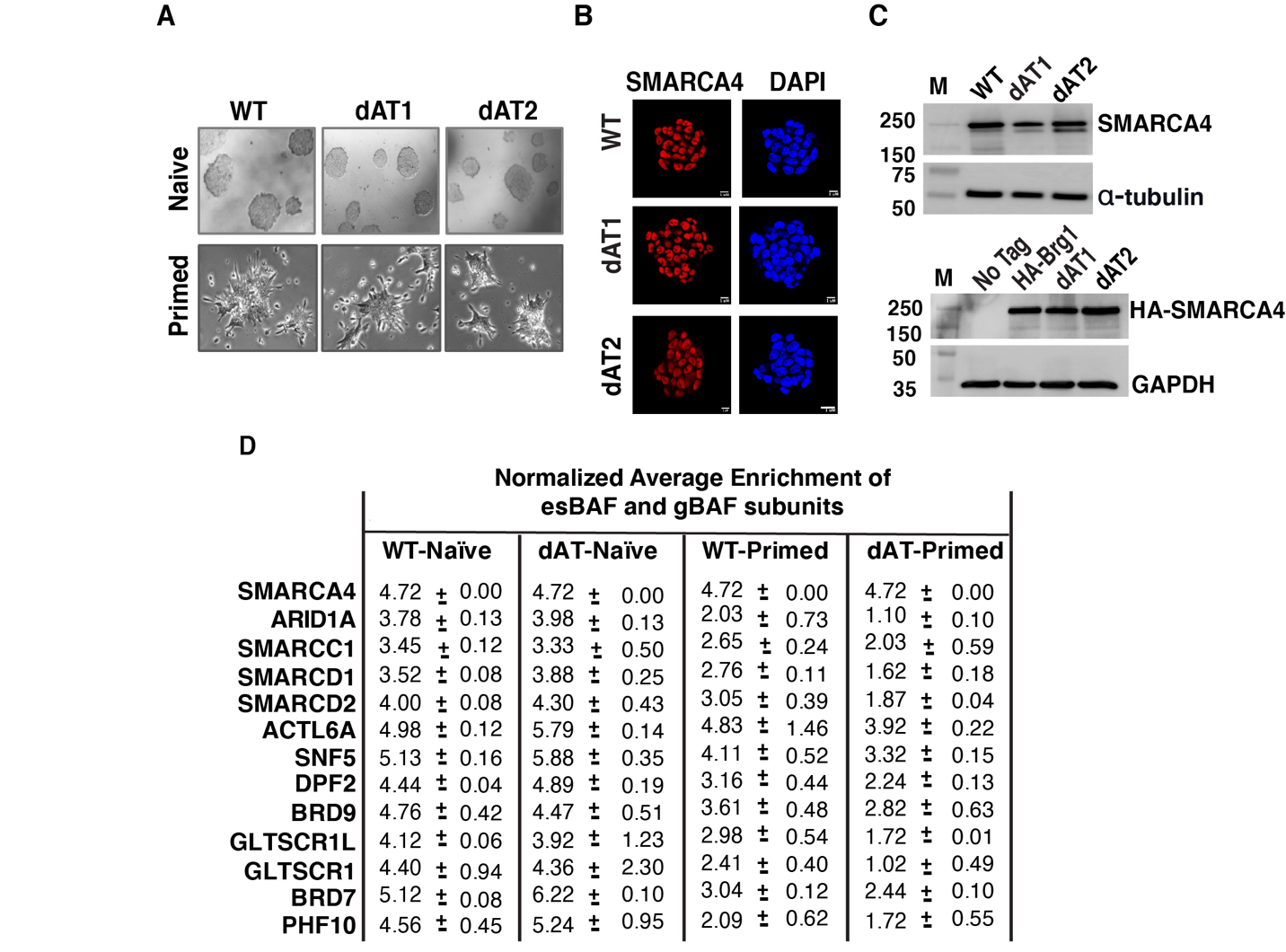

**Supplemental Figure S8. Complex integrity of esBAF is maintained after deletion of the AT-hook domain of SMARCA4.** (A**)** Morphology of wildtype (WT) and AT-hook deletion mutants (dAT1 and dAT2) mESCs grown in naïve [(LIF/2i media supplemented with leukemia inhibitory factor (LIF), MAP/ERK kinase inhibitor (MEKi), glycogen synthase kinase 3 beta inhibitor (GSK3βi)] and primed [(media supplemented activin A, fibroblast growth factor 2 (FGF2)] conditions. (B**)** The localization and expression of SMARCA4 in WT and dAT mutants is shown by immunofluorescence in naïve cells**.** (C**)** SMARCA4 expression in WT and AT-hook deletion mutants in naïve state is determined by immunoblotting (top), α-tubulin as a loading control. The bottom panel shows the western blot for HA-tagged SMARCA4 expression in WT and dAT mutant mESCs in naïve condition; untagged cells was used as a control and gapdh used as a loading control. (D**)** Table shows the average enrichment of es-BAF and gBAF components in WT and dAT mutant cells in both naïve and primed states.
